# Characterization of an amyloidogenic intermediate of transthyretin by NMR relaxation dispersion

**DOI:** 10.1101/2024.02.16.580707

**Authors:** Benjamin I. Leach, James A. Ferguson, Gareth Morgan, Xun Sun, Gerard Kroon, David Oyen, H. Jane Dyson, Peter E. Wright

**Author notes:** Address for correspondence: Peter E. Wright.

## Abstract

The aggregation pathway of transthyretin (TTR) proceeds through rate-limiting dissociation of the tetramer and partial misfolding of the monomers, which assemble into amyloid structures through a downhill polymerization mechanism. The structural features of the aggregation-prone monomeric intermediate are poorly understood. Characterization of amyloidogenic intermediates is challenging due to their propensity to aggregate at concentrations necessary for structural studies. NMR relaxation dispersion offers a unique opportunity to characterize these intermediates when they exchange on favorable timescales with NMR-visible ground states. To characterize the structural transitions associated with tetramer dissociation, we have analyzed ground-state chemical shift differences between the native tetramer and an engineered monomer in which the critical F87 side chain is replaced by glutamate. The secondary structure and overall fold of the F87E monomer is similar to that of the tetramer except for β-strand H. This strand populates two conformations, where it is either docked on the protein core or is displaced from the edge of the β-sheet formed by β-strands D, A, G, and H (DAGH β-sheet) and is dynamically disordered. Chemical shift differences derived from analysis of ^1^H/^15^N single, double and zero quantum relaxation dispersion data provide insights into the structure of a low-lying excited state that exchanges with the ground state of the F87E monomer at a rate of 3800 s^-1^. Disruption of the subunit interfaces of the TTR tetramer leads to destabilization of edge strands in both β-sheets of the F87E monomer. Conformational fluctuations are propagated through the entire hydrogen bonding network of the DAGH β-sheet, from the inner β-strand H, which forms the strong dimer interface in the TTR tetramer, to outer strand D which is unfolded in TTR fibrils. Fluctuations are also propagated from the AB loop in the weak dimer interface to the EF helix, which undergoes structural remodeling in fibrils. The conformational fluctuations in both regions are enhanced at acidic pH where amyloid formation is most favorable. The relaxation dispersion data provide insights into the conformational dynamics of the amyloidogenic state of monomeric TTR that predispose it for structural remodeling and progression to amyloid fibrils.

## Introduction

Transthyretin (TTR) is a highly-conserved transport protein in vertebrates, where it functions as a carrier for thyroxine T4 and retinol-binding protein.*^1^* TTR amyloidoses are progressive diseases that involve deposition of the protein as amyloid deposits in a variety of tissues.*^2,3^* In familial diseases such as familial amyloid cardiomyopathy (FAC) and familial amyloid polyneuropathy (FAP), inherited mutations destabilize the native tetrameric structure, resulting in dissociation and aggregation of TTR in the heart and peripheral nervous system, respectively. In elderly patients, senile systemic amyloidosis (SSA) results from deposition of wild-type TTR, a condition that affects up to 25% of the population over 80 years old.*^4^*

Transthyretin is one of a group of amyloidogenic globular proteins, which also includes superoxide dismutase (SOD-1), β2-microglobulin, immunoglobulin light chains, and lysozyme.*^5^* Aggregation of these globular amyloid precursors proceeds through formation of a partially misfolded intermediate, which transitions through various pre-amyloid structures before rearranging into mature fibrils.*^6^* The requirement for a partially misfolded intermediate in the aggregation pathway is thought to be a common feature for amyloidosis of globular proteins.*^7^* TTR is unique in this group of proteins in that it is assembled as a highly stable tetramer, with amyloidosis being limited by the rate of tetramer dissociation.*^6,8^*

TTR is assembled from four identical protomers, each composed of an eight-stranded β-sandwich (Figure 1). The β-sheets are formed from the DAGH and CBEF strands, with a single helix at the C-terminal end of strand E.*^9^* The interfaces of the tetramer are formed by antiparallel hydrogen bonding between the H-strands and, to a lesser extent, the F-strands of two protomers (the strong dimer interface), and through interactions between the AB and GH loops (the weak dimer interface). The strong dimer interface is also stabilized by packing of the sidechain of F87 into a pocket on the opposing protomer (Figure 1). The central channel between the DAGH sheets forms two T4 binding sites. As the hormone binding function of TTR is redundant with albumin and thyroxine binding globulin (TBG), the T4 binding sites are mostly unoccupied in the blood.*^10^* Ligand binding to the T4 sites stabilizes the tetrameric structure of TTR, a feature that has been exploited in the design of kinetic stabilizers such as tafamidis.*^11^*

**Figure 1.**
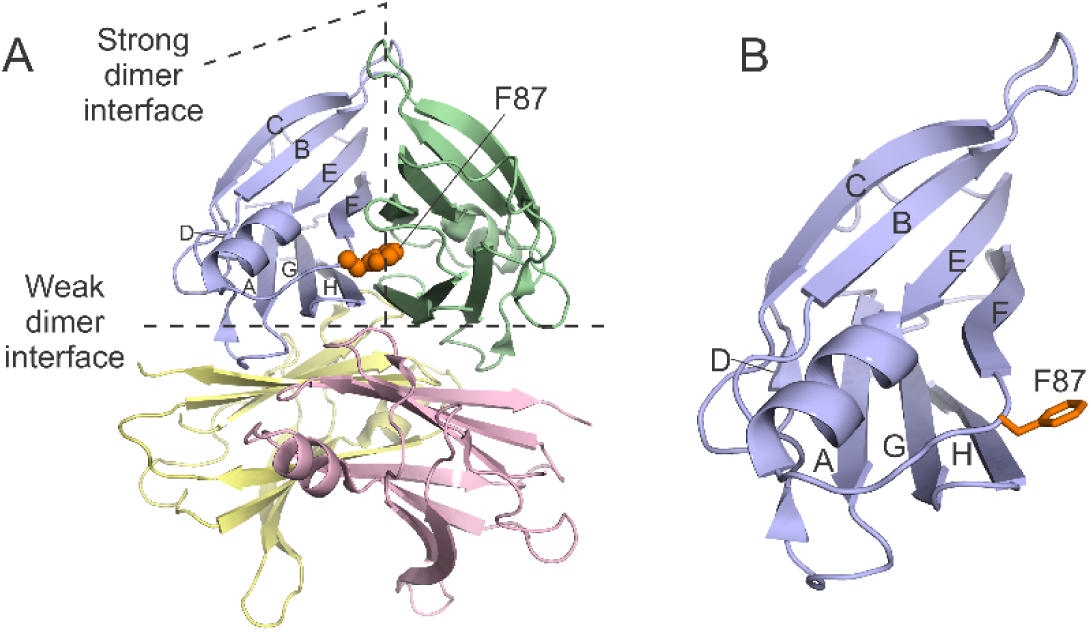
A. Cartoon representation of the native TTR tetramer (PDB code 5CN3)*^12^*. The strong and weak dimer interfaces are indicated by the hatched lines. B. Structure of the blue protomer from A. The F87 side chain, which was mutated to glutamate in F87E, is shown in orange.

The aggregation pathway of TTR was previously characterized by acid-induced denaturation experiments, which monitored native tryptophan fluorescence and tetramer dissociation as a function of pH. These studies showed that dissociation of the native tetramer occurs below pH 5 to form an aggregation-prone monomer with altered tertiary structure.*^6^* To assess the significance of this altered conformational state, Jiang *et al*.*^13^* engineered a monomeric form of TTR (M-TTR) by mutating F87 and L110 to methionine (F87M, L110M) to disrupt the strong and weak dimer interfaces, respectively. M-TTR is non-amyloidogenic at physiological pH, but forms fibrils when partially denatured by acidification. Biophysical and crystallographic studies showed that M-TTR at pH 7 is structurally very similar to the individual subunits in the intact TTR tetramer and partial unfolding at acidic pH is required to render it amyloidogenic.*^13^* NMR studies of M-TTR at physiological pH have revealed conformational fluctuations that release β-strand H and result in transient formation of a small population of an aggregation-prone state.*^14,15^* These fluctuations are suppressed by the stabilizing T119M mutation.*^14,16^*

Misfolded amyloidogenic intermediates remain poorly characterized, due to their short lifetime and tendency to aggregate. These intermediates are of great interest, since an understanding of the mechanism by which they assemble into amyloid is likely to provide insights into novel therapeutic approaches.*^17,18^* Since these intermediates are thermodynamically accessible from the ground state,*^5^* they may be studied using NMR Carr-Purcell-Meiboom-Gill (CPMG) relaxation dispersion methods,*^19^* provided conditions can be found where a small population of the amyloidogenic conformation exchanges with the ground state on a millisecond to microsecond timescale. The combined analysis of chemical shift data from multi-probe relaxation dispersion measurements provides a sensitive and powerful method for characterization of dynamic conformational changes in proteins. *^19,20^* For example, both ^15^N chemical shifts and ^13^C’ chemical shifts are sensitive to changes in backbone dihedral angles, but ^15^N chemical shifts are also extremely sensitive to changes in hydrogen bonding and therefore provide complementary information.*^21^* Similarly, ^1^H chemical shifts are sensitive to hydrogen bonding, but are also strongly affected by ring currents, local charge, and other effects.*^22^*

^15^N relaxation dispersion measurements at neutral pH identified regions of M-TTR where structural fluctuations on a μs time scale lead to formation of a small and transient population of an aggregation-prone excited state.*^14^* While these studies provided important insights into the molecular basis of monomer unfolding, neither the kinetics of the process nor the conformation of the partially unfolded excited state could be ascertained using ^15^N relaxation dispersion data alone. In the present work we extended the relaxation dispersion measurements to additional nuclei to better define the kinetics and structural changes associated with ground state conformational fluctuations in the TTR monomer. Rather than using M-TTR, where the double mutation F87M/L110M destabilizes both the strong and weak dimer interfaces,*^13^* we engineered a new single-site monomeric mutant of TTR (F87E) with improved qualities for NMR structural studies and relaxation experiments. Here we compare the solution structures of the WT TTR tetramer and the folded F87E monomer based on backbone chemical shifts, and report amide ^1^H and ^15^N chemical shift changes for the amyloidogenic intermediate determined from a suite of ^1^H_N_ and ^15^N_H_ single, double and zero quantum relaxation dispersion experiments. We have also performed ^13^C’ single quantum relaxation dispersion experiments, which are primarily sensitive to changes in backbone dihedral angles and are complementary to the amide dispersion data.

Together, these chemical shifts reveal detailed conformational changes that correlate well with previously observed features of the amyloidogenic state and suggest a mechanism for the coupling between tetramer dissociation and monomer misfolding.

## Results

### Engineering a monomeric TTR for relaxation dispersion studies

For the relaxation dispersion experiments, we introduced a single mutation, F87E, reasoning that the negatively charged glutamate residue would be more unfavorable than the methionine of M-TTR for packing into the hydrophobic F87 binding pocket in the neighboring subunit across the strong dimer interface (Figure 1A). To verify that this mutant was completely monomeric, we conducted pulsed-field gradient spin-echo diffusion experiments*^23–25^* on F87E at two concentrations, 150 and 300 μM. The results of these experiments are shown in Figure S1A in the Supplementary Material, and yield almost identical diffusion constants, (7.0 ± 0.1) x 10^-7^ and (6.9 ± 0.08) x 10^-7^ cm^2^s^-1^ at 150 μM and 300 μM, respectively. ^19^F NMR spectra of a TTR mutant (C10S/S85C/F87E), labeled at position 85 with 3-bromo-1,1,1-trifluoroacetone (BTFA) show that the F87E mutant remains monomeric up to at least 470 μM concentration (Figure S1B). F87E remains monomeric even in the presence of tafamidis, a ligand that stabilizes the TTR tetramer*^11^* (Figure S1C). To ensure that the F87E monomer does not transiently self-associate, which would complicate analysis of the relaxation dispersion data, we conducted ^1^H_N_ single-quantum relaxation dispersion experiments at 150 and 300 μM concentration. The dispersion curves are the same, within experimental error, at the two concentrations (Figure S1C). We conclude that the F87E mutation is sufficient to completely disrupt tetramer assembly under the conditions used for relaxation dispersion measurements.

The aggregation behavior of F87E was compared with that of wild-type TTR and M-TTR (Figure S2). At pH 4.3 and 37 °C, both F87E and M-TTR readily form thioflavin T-binding aggregates over four hours, with indistinguishable kinetics (Figure S2A). Under the same conditions, wild-type TTR did not form aggregates. None of the variants had aggregated at pH 7 after 24 h. To verify that the TTR variants remained soluble, we measured absorbance at 330 nm, which we attribute to light scattering by aggregated TTR, before and after the aggregation time course *^6,26^* (Figure S2B). Consistent with the thioflavin T kinetics, the turbidity of both F87E and M-TTR solutions increased to a greater extent than that of wild-type TTR after 24 h. These data showed that the aggregation behavior of F87E is similar to M-TTR, showing no aggregation at physiological pH but forming amyloid more rapidly than wild-type TTR under partially denaturing conditions at pH 4.3.*^13^*

### Assignment of F87E NMR spectra

As with M-TTR, ^1^H,^15^N HSQC spectra of F87E showed severe broadening for several cross peaks and variable peak intensity due to the inherent conformational dynamics of the protein (Figure 2, full spectrum in Figure S3). In order to assign the resonances of F87E, we employed a strategy of sparse sampling and co-processing of triple resonance spectra using coMDD.*^27,28^* We were able to assign amide cross peaks for 103 of 119 non-proline residues, including many backbone resonances for the DE loop, the helix, FG loop, β-strand H, and the C-terminal region that were not observable in the spectra of M-TTR at similar pH.*^14^* We also observed several broad and weak cross peaks in the HSQC spectrum of F87E that we were unable to assign because they lacked sequential connectivities in triple resonance spectra.

**Figure 2.**
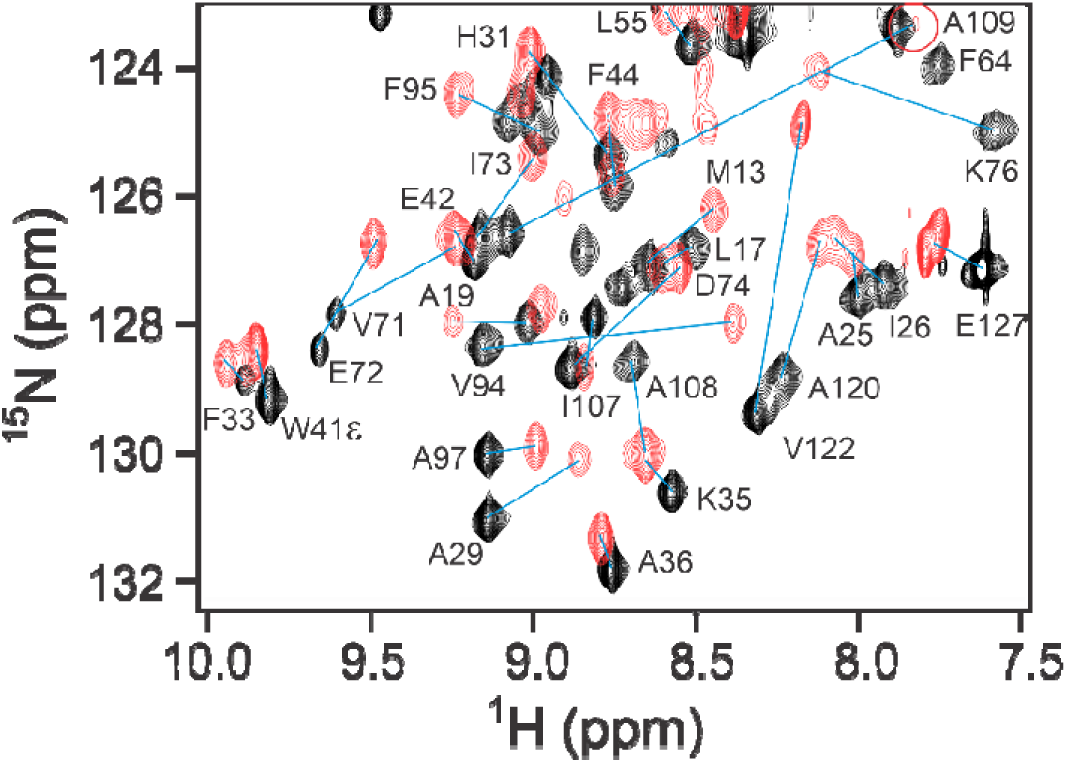
Region of the ^1^H,^15^N HSQC spectrum of tetrameric WT TTR (black), superimposed on the same region of F87E TTR (red) at pH 6.5 and 298K. Selected assignments are shown and corresponding cross peaks in the two spectra are joined by a blue line. The weak cross peak of A109 in the F87E spectrum is shown with a red circle.

### Conformational differences between monomeric and tetrameric TTR

M-TTR was originally engineered to determine the importance of partial misfolding of the TTR monomer for entry into the amyloid cascade. Dissociation of the native tetramer and misfolding of the monomer are strongly coupled thermodynamically,*^29^* and occur simultaneously under partially denaturing conditions.*^6^* Accounting for structural differences between the protomer in the TTR tetramer and the ground state of the TTR monomer is therefore essential for meaningful analysis of structural changes between the folded and misfolded monomer. NMR structures have been reported for M-TTR at high pressure (500 bar), where the spectral quality is improved.*^15^* The overall structure of M-TTR is similar to that of the protomer in the wild-type TTR tetramer, except that the H strand is dissociated and there are conformational changes in the AB and FG loops. Although the quality of TTR F87E spectra at atmospheric pressure is substantially better than for those of M-TTR, structure determination still presents a major challenge due to exchange broadening of a significant number of resonances. In particular, the amide ^1^H-^15^N cross peaks of residues 87-92 (EF loop and N-terminus of F strand) and 109-117 (GH loop, C- and N-terminal regions of G and H strands, respectively) are broadened and/or shifted from their positions in the HSQC spectrum of the wild-type TTR tetramer (Figure S3) and could not be assigned. We therefore accounted for the structural differences between the ground state of monomeric TTR F87E and the protomer in tetrameric TTR by comparing chemical shifts (Figure 3A) and φ and ψ dihedral angles predicted using TALOS-N.*^30^*

**Figure 3.**
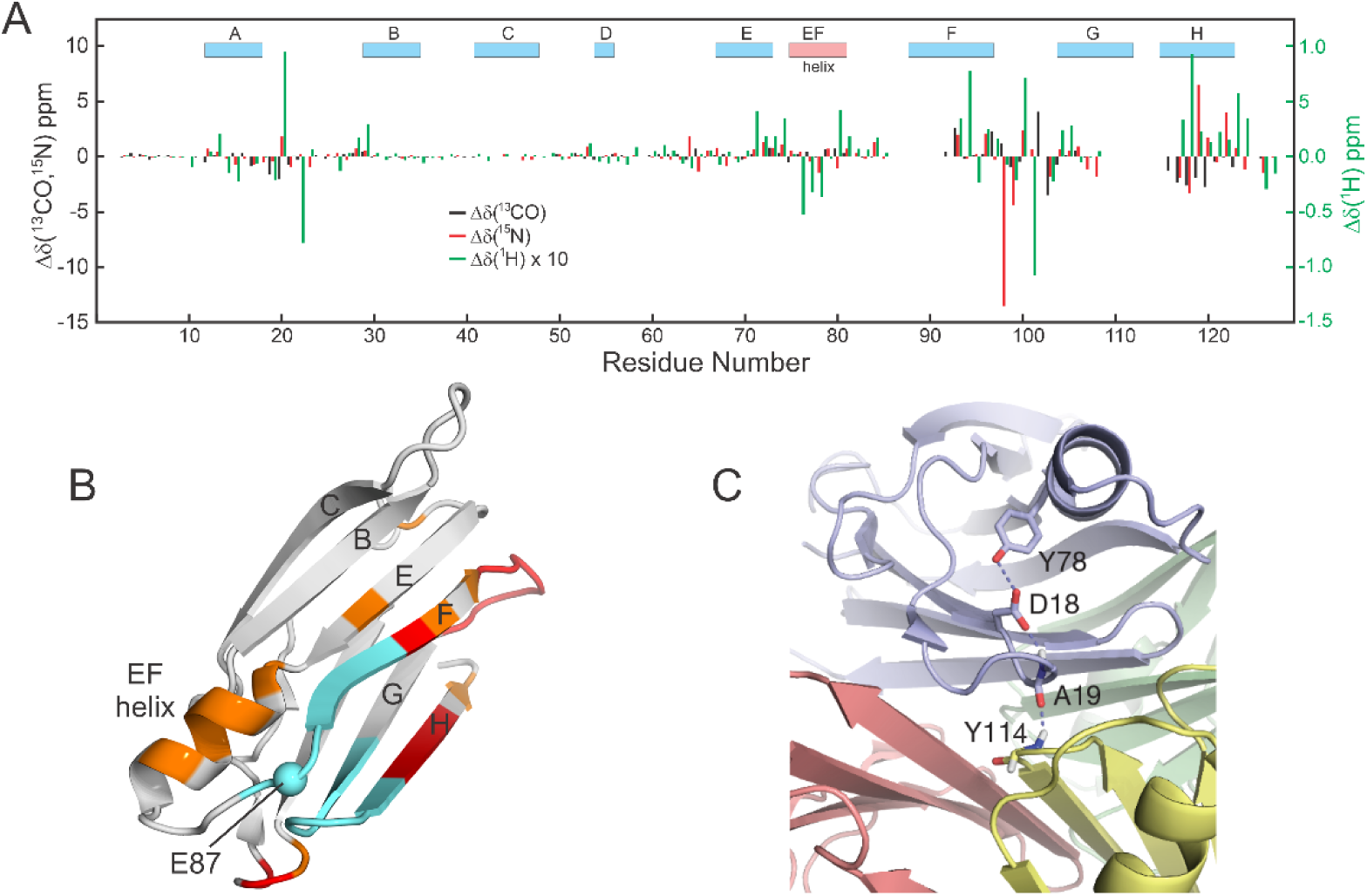
A. Chemical shift differences between WT-TTR and F87E (δ_WT_– δ_F87E_) at pH 6.5, 298K. The colored rectangles at the top of the figure indicate the locations of secondary structure elements in the X-ray crystal structure, β-strands A-H (blue) and the EF-helix (pink). B. Cartoon structure of the TTR protomer (PDB 5CN3) showing weighted average ^1^H_N_, ^15^N, ^13^C’ chemical shift differences (Δδ_ave_) between F87E and the WT tetramer. Residues with shift differences greater than 2 standard deviations (red) and 1-2 standard deviations (orange) are indicated. Regions where amide resonances were not observed are shown in cyan. Chemical shifts were scaled using the standard deviations of the ^1^H_N_, ^15^N, ^13^C’ chemical shifts in the Biological Magnetic Resonance Databank. The weighted average shift difference was calculated as Δδ_ave_ = (Δδ_H_^2^ + (Δδ_N_/6.27)^2^ + (Δδ_C’_/2.98)^2^)^1/2^ C. Region of the TTR tetramer showing the inter-protomer hydrogen bond between Y114 and A19 that is absent in the monomer.

TTR commonly crystallizes in space group P2_1_2_1_2 with two protomers in the asymmetric unit. Analysis of the wild type TTR tetramer chemical shifts using TALOS-N*^30^* confirms that the secondary structure observed in the crystal is closely maintained in solution. Interestingly, the conformation of the FG loop in solution is close to that of chain A in the P2_1_2_1_2 crystal structures and differs substantially from that of chain B (Figure S4).

Chemical shift differences between TTR F87E and the wild-type TTR tetramer are observed in the AB loop, EF helix, F β-strand, FG loop and the C-terminal region of the H β-strand (Figure 3A). Weighted average ^1^H_N_, ^15^N, ^13^C’ chemical shift differences are plotted onto the protomer structure in Figure 3B. Most of the chemical shift differences reflect the disruption of the strong and weak dimer interfaces and the associated loss of hydrogen bonds. In the AB loop, large chemical shift changes are observed due to the disruption of the hydrogen bond network that stabilizes the weak dimer interface. Most notably, upon tetramer dissociation the hydrogen bond between A19CO and Y114NH (Figure 3C) is broken, leading to a large 2.2 ppm change in the V20 ^15^N chemical shift. The V20 ^1^H_N_ resonance shifts 0.88 ppm upfield in F87E, likely reflecting disruption of the V20 NH – D18 O^δ^ hydrogen bond upon dissociation of the tetramer. These residues form part of a hydrogen bond network that connects the EF helix to the weak dimer interface through the sidechains of Y78 and D18 (Figure 3C). The G22 ^1^H_N_ resonance shifts 0.86 ppm downfield in the F87E spectrum due to loss of the ring current from Y114 in the neighboring subunit across the weak dimer interface. ^13^C’ chemical shifts and backbone dihedral angles predicted by TALOS-N (see later) indicate conformational changes in the AB loop associated with disruption of the weak dimer interface.

Residues in the strong dimer interface also exhibit substantial chemical shift changes between the F87E monomer and the wild-type tetramer. Large ^13^C’ and ^15^N chemical shift changes are observed for residues 117-123 in the H strand, reflecting the loss of H-H’ hydrogen bonds across the strong dimer interface and perturbation of intra-protomer G-H hydrogen bonding. In F87E, the chemical shifts of residues 119 -126 are close to sequence-corrected random coil values,*^31^* showing that strand H is flexible from residue 119 to the C-terminus. Amide resonances of residues in the EF loop, near the site of the F87E substitution, are broadened beyond detection, as are resonances from the N-terminal part of strand F. Large ^1^H_N_ and ^15^N chemical shift changes are observed for residues at the C-terminal end of strand F due to loss of inter-protomer hydrogen bonds. The disruption of the strong dimer interface results in structural changes to the FG loop. The strong dimer interface is stabilized by packing of the aromatic sidechain of F87 into a hydrophobic pocket between the F’ and H’ strands in the opposing protomer, where it packs against the aromatic rings of F95 and Y105. The magnitude of the ^13^C’, ^15^N, and ^1^H_N_ chemical shift changes indicates a substantial change in the conformation and hydrogen bonding pattern of the FG loop, likely associated with removal of the F87 ring from its binding pocket upon formation of the monomer. A conformational rearrangement of the FG loop was also observed in the solution structure of M-TTR.*^12,15^*

Backbone φ and ψ dihedral angles and chemical shift order parameters for F87E and the WT TTR tetramer were predicted from ^15^N_H_, ^1^H_N_, ^13^CO,^13^Cα and, where available, ^13^Cβ chemical shifts using TALOS-N.*^30^* Results are shown in Figure S5A. The predicted dihedral angles of F87E are overall very similar to those predicted for the WT TTR tetramer except for small differences in the AB loop, C and D strands, DE, and FG loops, and the C-terminal region of the H strand. The largest conformational changes occur in the FG loop; the ^15^N resonance of N98 is shifted 13.1 ppm downfield in the spectrum of F87E, reflecting a change in backbone conformation from the α_R_ region in the tetramer to the polyproline II region in F87E. Chemical shift order parameters*^32^* calculated using TALOS-N show increased flexibility relative to the WT tetramer in the N-terminal region, the AB and EF loops, near the site of the F87E substitution, and in the C-terminal region of the H strand (Figure S5B).

### The C-terminal region of the H strand of F87E adopts two conformations

As noted above, there are several broad and weak cross peaks in the ^1^H,^15^N HSQC spectrum of F87E that lack sequential connectivities in triple resonance spectra. Two of these cross peaks (at 8.50, 125.1 ppm and 8.46, 120.8 ppm, Figure S6A) occur at very similar chemical shifts to the cross peaks of A120 (8.52, 125.1 ppm) and V121 (8.45, 121.0 ppm) in spectra of M-TTR (Figure S6B).*^14^* The amide at 8.50 and 125.1 ppm is connected to a ^13^C^β^ resonance at 18.7 ppm in an HNCB spectrum of F87E and to a ^1^H resonance at 1.13 ppm in a ^1^H-^15^N TOCSY spectrum, identifying it as the amide of an alanine (Figure S7). Both the 8.50 and 8.46 ppm amide ^1^H resonances exhibit NOEs to the 1.13 ppm methyl resonance in a ^1^H-^15^N NOESY spectrum of F87E (Figure S7), consistent with assignment to A120 and V121. However, in common with M-TTR,*^14,15^* assignments could not be extended to neighboring residues. A second set of amide cross peaks for A120 and V121, which could be assigned unambiguously from triple resonance spectra in which there are unambiguous connectivities linking residues S117 through N124, were identified at near random coil chemical shifts (8.16, 126.9 ppm and 8.02, 120.2 ppm, respectively), as described above. These cross peaks as well as those of T119, V122, and T123 (labeled in blue in Figure S6) are absent from the HSQC spectrum of M-TTR (Figure S6B).*^14,15^* The C-terminal E127 cross peak is also split in the HSQC spectrum of F87E, but not that of M-TTR, with components of approximately equal intensity at 7.83, 127.1 ppm and 7.79, 127.0 ppm. Thus, there appear to be two states of F87E in which the H strand adopts alternative conformations that are in slow exchange on the NMR timescale, one with intense cross peaks at approximately random coil chemical shifts and the other with broad and weak HSQC cross peaks. We refer to these as state A, in which the chemical shifts of T119 to E127 are close to random coil values, and state B where chemical shifts of A120 and V121 are close to M-TTR values and residues 119 and 122-124 could not be assigned. In state A, the A120 and V121 cross peaks exhibit large negative amide proton chemical shift temperature coefficients (Δδ/ΔT = −5.9 and −9.0 ppb/K for A120 and V121, respectively), indicating solvent exposure and the absence of intramolecular hydrogen bonds.*^33,34^* In contrast, in state B, Δδ/ΔT for the A120 and V121 resonances is only weakly negative (−1.2 and −2.0 ppb/K, respectively), consistent with a conformation in which the C-terminal region of the H strand is docked and participates in hydrogen-bonded secondary structure. Exchange between states A and B is slow and the bound state peaks are very weak; attempts to directly observe exchange between the states using chemical exchange saturation transfer (CEST) and ZZ-exchange experiments were unsuccessful.

### Multi-nuclear relaxation dispersion measurements characterize conformational exchange in F87E

Similar to M-TTR,*^14^* the F87E monomeric TTR variant undergoes conformational exchange that leads to NMR relaxation dispersion. To characterize the exchange process, we performed ^1^H/^15^N single, double and zero quantum CPMG experiments, which when fitted together improve the accuracy and robustness of the kinetic parameters and the chemical shift differences between the ground and excited states.*^35^* ^1^H_N_ and ^15^N_H_ single, double and zero quantum relaxation dispersion profiles recorded on 500 and 800 MHz spectrometers were fitted to the Carver-Richards equation*^36^* using the software package GLOVE.*^37^* The data were fit to a two-site exchange model (A ↔ B) to derive a global exchange rate constant *k_ex_*, the populations *p_A_* and *p_B_*, the forward and backward rate constants *k_A_* and *k_B_*, and the single quantum (ΔωH and ΔωN), zero quantum (Δω_ZQ_ = ΔωH – ΔωN), and double quantum (Δω_DQ_ = ΔωH + ΔωN) chemical shift differences between the exchanging states. Representative dispersion profiles and their fits are shown in Figure 4 and fitted dispersion profiles for all residues are shown in Figure S8. Global values of *k_ex_* (3800 ± 80 s^-1^) and *p_B_* (5.4 ± 0.3 %) were determined by fitting data for 15 residues with well-defined dispersion curves and minimal scatter in the data points; uncertainties were estimated from 200 Monte Carlo simulations. The exchange rate is relatively fast on the CPMG timescale, and the accuracy of the parameters derived from fits to single quantum dispersion data alone is limited due to correlation between *k_ex_* and *p_B_* at fast exchange rates. The inclusion of the double and zero quantum relaxation dispersion data in the analysis improves the accuracy of the fitted parameters. The remaining relaxation dispersion profiles were fitted to the Carver-Richards equation using the global *k_ex_* and *p_B_* values determined from the initial set. For the majority of assigned residues, the ^1^H, ^15^N SQ/DQ/ZQ dispersion profiles fit well to a two-site exchange model.

**Figure 4.**
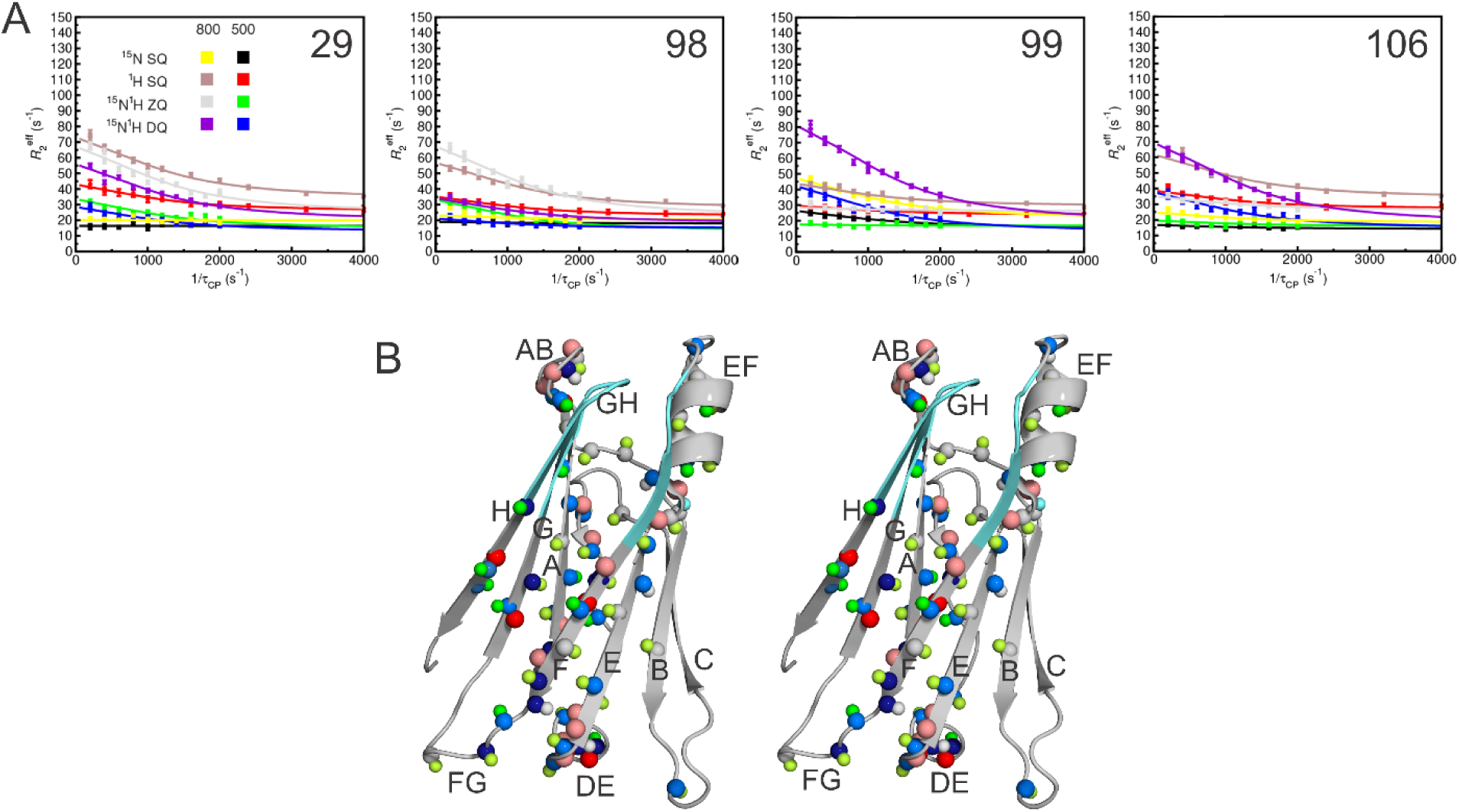
A. Representative relaxation dispersion profiles at 800 MHz and 500 MHz, fitted simultaneously to global exchange parameters (*k_ex_*and populations *p_A_* and *p_B_*) using GLOVE.*^37^* The dispersion profiles are color coded as: red, 500 MHz ^1^H single-quantum (SQ); brown, 800 MHz ^1^H SQ; black, 500 MHz ^15^N SQ; yellow, 800 MHz ^15^N SQ; green, 500 MHz ^1^H - ^15^N zero-quantum (ZQ); gray: 800 MHz ^1^H - ^15^N ZQ; blue, 500 MHz ^1^H - ^15^N double-quantum (DQ); purple, 800 MHz ^1^H - ^15^N DQ. The solid lines represent fits to the Carver-Richards equation*^36^* with fixed *k_ex_* (= 3800 s^-1^) and *p_B_*(= 0.054) derived from fitting the ^15^N and ^1^H_N_ SQ, ZQ, and DQ dispersion data for a cluster of 15 residues with well-defined dispersion curves and minimal scatter in the data points. B. Stereo view of a carton structure of the TTR protomer (from PDB entry 5CN3)*^12^*. Spheres indicate the location of residues that exhibit relaxation dispersion for ^15^N, ^1^H_N_, or ^13^C’ resonances. The spheres are color coded as follows: dark blue, ΔωN > 1.5 ppm; light blue, 0.5 < ΔωN < 1.5 ppm; dark green, ΔωH > 0.25 ppm; light green, 0.1 < ΔωH < 0.25 ppm; red, ΔωC’ > 1.0 ppm; pink, 0.5 < ΔωC’ < 1.0 ppm. Regions where amide resonances were not observed are shown in cyan.

The sign of Δω, the chemical shift difference between the ground and excited states, cannot be determined from the relaxation dispersion data. However, under favorable conditions, differences in the peak positions in ^1^H, ^15^N HSQC and HMQC spectra can provide signs for the ^15^N chemical shifts, with signs for the ^1^H_N_ chemical shifts determined indirectly using the relative signs available from the DQ and ZQ relaxation dispersion data.*^35,38^* Following the criteria of Skrynnikov *et al*.,*^38^* we determined the sign of Δω_N_for a subset of residues for which the exchange-induced shift between HSQC and HMQC spectra Ω_N_ is > ∼0.3 Hz, and where |ξ_N_| (= |Δω_N_|/ k_B_) is > ∼0.04 Hz and |ξ_H_| (= |Δω_H_|/ k_B_) is comparable to or greater than |ξ_N_|. Residues with severely overlapped cross peaks were excluded from the analysis. The values of Δω derived from the relaxation dispersion data are summarized in Table S1. The available signs provide valuable information about structural changes in the excited state of the monomeric TTR.

To supplement the amide dispersion data, we performed ^13^C’ relaxation dispersion experiments on uniformly ^13^C-^15^N labeled F87E using the approach of Lundström *et al. ^39^* The relaxation dispersion profiles are shown in Figure S9, for residues that exhibit ^13^C’ relaxation dispersion. Chemical shift differences (Table S2) were determined by fitting the data to the global *k_ex_* and *pB* obtained from the combined fits of the ^1^H and ^15^N data. Carbonyl chemical shifts are primarily influenced by primary structure and backbone and sidechain dihedral angles and are largely insensitive to hydrogen bonding,*^22^* so are complementary to amide dispersion for interpreting structural changes. Due to the fast exchange rate and the relatively small values of ΔωC’, the ^13^C’ dispersion curves are largely featureless, but are still valuable to complement the analysis of the amide data.

Previous ^15^N relaxation dispersion studies on M-TTR revealed fluctuations across the DAGH sheet and in the AB and EF loops.*^14^* Our data for F87E show a similar pattern of dispersion as the previous study of M-TTR, but the increased assignment coverage, improved spectral quality of F87E, and acquisition of SQ, ZQ, and MQ relaxation dispersion profiles for ^1^H and ^15^N enabled a more complete description of the dynamic behavior of the TTR monomer. We also observed ^13^C’ dispersion for 23 residues, but the carbonyls of only four residues (D18, E62, V65, V93, Y105, and T119 in state B) have ΔωC’ greater than 1 ppm, implying little change in backbone dihedral angles in the excited state. It is notable that the residues that exhibit ^13^C’ dispersion are mostly located in the AB and DE loops, the D strand, the EF helix, and strand F (Figure 4B). Many residues in the DAGH and CBEF β-sheets exhibit ^1^H_N_ and ^15^N dispersion with little observable ^13^C’ dispersion, suggesting propagation of structural perturbations across the sheet through the hydrogen bond network with little change in backbone conformation. The location of the dispersing residues is mapped onto the TTR protomer structure in Figure 4B.

### F87E adopts two ground state conformations that differ in dynamics

For state A of F87E, residues 120-127 at the C-terminus of the H strand have small R_2_ values and exhibit no relaxation dispersion (Figure S8), consistent with their near random coil chemical shifts and fast time scale dynamics. In contrast, the resonances of A120 and V121 in state B exhibit strong ^1^H_N_ and ^15^N dispersion (Figure S8), implying fluctuations in their hydrogen bonding interactions, most likely with residues in strand G. In the native protomer structure, a pair of hydrogen bonds is formed between the NH and carbonyl groups of V121 and T106, which also exhibits strong relaxation dispersion (Figure 4, S8). Resonances of residues 122-126 could not be assigned for state B but one component of the split E127 cross peak (7.79, 127.0 ppm) exhibits weak dispersion. Residues 120, 121, and 127 also exhibit ^15^N relaxation dispersion in M-TTR.*^14^*

### Nature of the F87E excited state

Mutant monomeric TTR is stable near neutral pH but aggregates rapidly when partially unfolded at pH 4.4. *^13,14,40^* We have previously reported pH titrations of F87E, performed at 277K to decrease the rate of aggregation.*^40^* As the pH is reduced from 6.7 to 4.4, amide cross peaks associated with residues in strand A, the AB loop, strand D, the DE loop, the EF loop, and the C-terminal end of strand F lose more than 60% intensity. All of these residues exhibit relaxation dispersion at pH 6.5 (Table S1, Figure S8) and the intensity loss is consistent with enhanced conformational fluctuations at pH 4.4 that lead to exchange broadening.*^40^* In contrast, amide resonances of residues in the EF helix and strand H become more intense, indicating that these regions become more flexible at acidic pH, suggesting local unfolding. Further analysis of the titration data reveals that the A120 and V121 cross peaks in state B lose intensity as the pH is reduced from 6.7 to 4.4, with concomitant increase in the intensity of the corresponding cross peaks in state A (Figure S10). It thus appears that under the acidic conditions that favor aggregation, the strand G – strand H interactions in state B are disrupted to free the C-terminal region of the H strand and increase the population of state A. The correlation between residues showing relaxation dispersion and those undergoing exchange broadening at acidic pH (Table S1) strongly suggests that the excited state sampled through conformational fluctuations of F87E near neutral pH corresponds to the aggregation-prone amyloidogenic state. Similar observations were made for M-TTR.*^14^*

### Perturbation of hydrogen bonding networks in the amyloidogenic excited state

In the TTR tetramer, the aromatic ring of F87 extends across the strong dimer interface and packs into a hydrophobic pocket formed by V93, F95, Y105, I107, A120, and V122 of the neighboring protomer. V94, F95, and I107 show ^1^H_N_ and ^15^N dispersion in the F87E monomer and the V93 and Y105 carbonyls exhibit strong ^13^C dispersion with ΔωC’ 2.4 and 1.2 ppm, respectively, reflecting a structural rearrangement in the F-strand and at the N-terminus of the G strand upon removal of the F87 ring. The amide resonances of R104 and Y105 are exchange broadened to the extent that we were unable to extract meaningful dispersion data.

The ^1^H_N_ and ^15^N dispersion observed for monomeric F87E TTR can be attributed to changes in backbone hydrogen bonding interactions between the ground and excited states, The amide groups that exhibit ^15^N and/or ^1^H_N_ dispersion and their associated hydrogen bonding networks in the native TTR protomer (chain B of the tetramer structure 5CN3) are shown in Figures 5A-D. ^15^N chemical shifts are exquisitely sensitive to hydrogen bond length; changes in hydrogen bonding to a peptide carbonyl group lead to large changes (up to 4.5 ppm) in the amide

**Figure 5.**
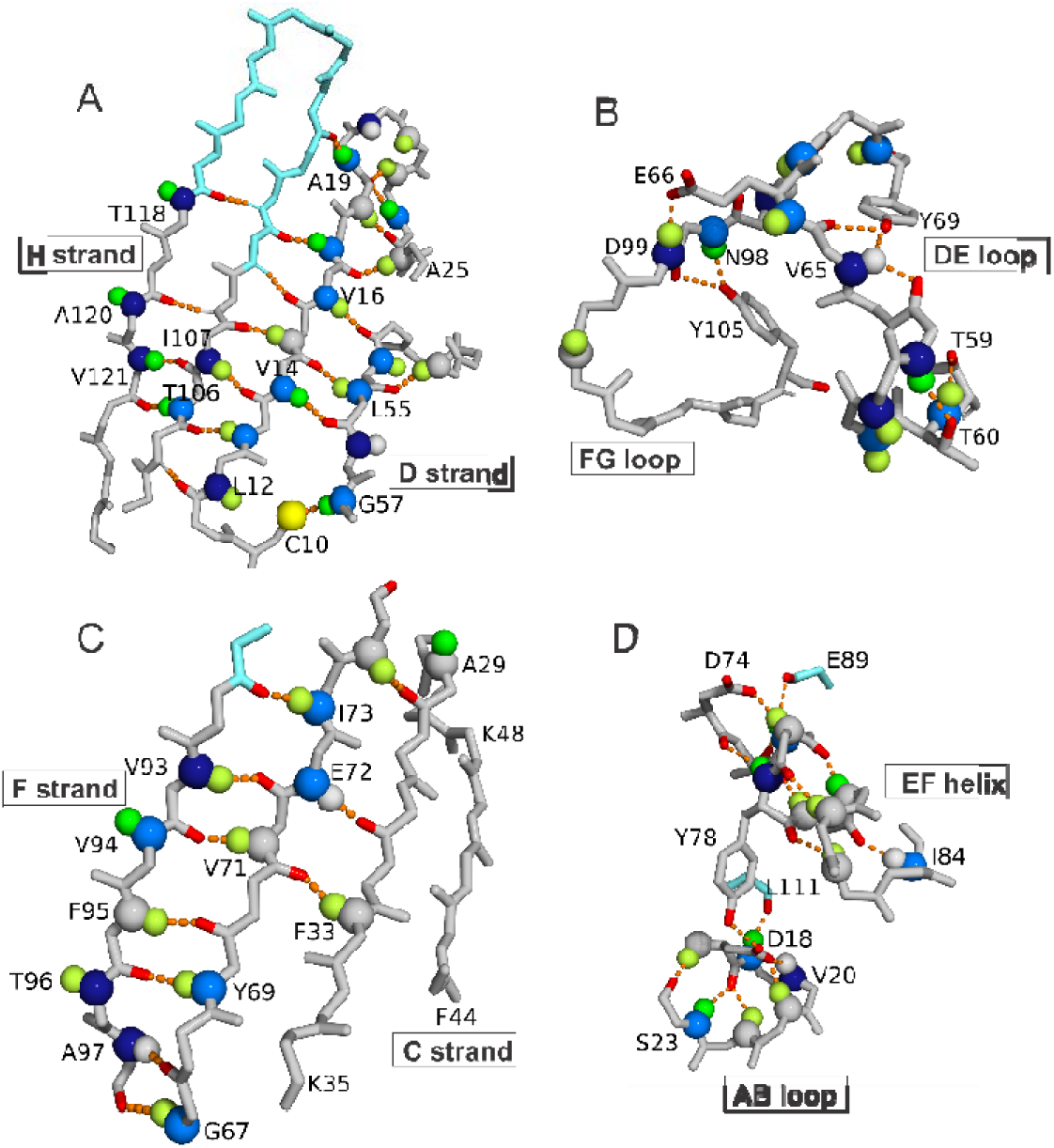
Fluctuations in hydrogen bonding networks of F87E monomer identified from analysis of relaxation dispersion data. A. DAGH β-sheet. B. DE and FG loops. C. CBEF β-sheet. D. AB loop and EF helix. The figures were made from the coordinates of PDB entry 5CN3. Hydrogen bonds inferred from the X-ray structure are depicted by hatched orange lines. Spheres indicate the location of residues that exhibit relaxation dispersion for ^15^N, ^1^H_N_ resonances. The spheres are color coded as follows: dark blue, ΔωN > 1.5 ppm; light blue, 0.5 < ΔωN < 1.5 ppm; dark green, ΔωH > 0.25 ppm; light green, 0.1 < ΔωH < 0.25 ppm. Residues whose amide resonances were not observed are shown in cyan.

^15^N shift whereas changes in direct hydrogen bonding to the NH group cause changes in both ^15^N and ^1^H_N_ chemical shifts.*^21^* Fluctuations are observed in the hydrogen bonding network across the entire DAGH sheet (Figure 5A); resonances of residues 109-117 that form the GH hairpin could not be assigned and appear to be severely broadened by the exchange process. Large ^15^N chemical shift changes are observed for residues in strands G (I107) and H (A120 and V121 in state B, T118) indicating substantial perturbations in interstrand hydrogen bonding. ^13^C’ dispersion is observed for the carbonyl groups of Y105 and T119 (in state B), suggesting changes in backbone conformation in the excited state. Large ΔωN values (> 1.8 ppm) are also observed for L12 and H56, pointing to substantial perturbation of the hydrogen bonding interactions between the N-terminus of strand A and the short D strand (residues 54-56). The amide resonances of V16 shift upfield (ΔωN = −0.57ppm, ΔωH = −0.16 ppm), suggesting a possible weakening of the hydrogen bond to the G53 carbonyl in the excited state. The amide resonances of G57 shift downfield (ΔωN = 1.08 ppm, ΔωH = 0.29 ppm) towards random coil positions predicted using POTENCI*^41^* (ΔδN = 1.46 ppm, ΔδH = 0.13 ppm), likely reflecting weakening or loss of the C10 Sγ – G57 NH hydrogen bond observed in TTR X-ray structures. The observation of ^13^C’ dispersion for the carbonyl groups of P11, L12, G53, and E54 provides further indication of perturbed interactions between strands A and D in the excited state.

The structural perturbations in the D strand of the excited state propagate into the DE loop (residues 57-67). ^13^C’ dispersion is observed for residues 61-63 and 66-68, indicating backbone dihedral angle changes in this region. ^15^N and ^1^H_N_ dispersion reveals accompanying perturbations in hydrogen bonding, both in the DE loop and neighboring FG loop. The hydrogen bonding in these regions inferred from chain B of the X-ray structure 5CN3, where the sign of the L dihedral angle of N98 is negative in accord with the solution data for F87E, is depicted in Figure 5B. In the X-ray structure, there are extensive hydrogen bonding interactions between backbone and side chains. While comparison of backbone dihedral angles predicted from chemical shifts for F87E with those in the X-ray structure 5CN3 suggests small differences in the conformation of these loops, it is of note that fluctuations of the side chain-backbone hydrogen bonds inferred from the crystal structure would be consistent with the observed pattern of relaxation dispersion. Within the DE loop of the X-ray structure, the hydroxyl groups of T59 and T60 are hydrogen-bonded to the amide NH of T60 and E63, respectively, while the Y69 hydroxyl potentially forms hydrogen bonds to V65 NH and carbonyl oxygen. The amide resonances of these peptide groups exhibit ^15^N and ^1^H_N_ relaxation dispersion. In the FG loop, the ^15^N and ^1^H_N_ resonances of N98 and D99 disperse strongly, likely as a result of fluctuations in hydrogen bonds between the Y105 hydroxyl group and the NH and CO moieties of N98. Relaxation dispersion is also observed for E61 and E62 in the DE loop; this cannot be explained by the static X-ray structures and likely reflects local conformational fluctuations that lead to altered hydrogen bonding patterns.

Fluctuations in the hydrogen bonding network in the CBEF sheet are less pervasive than for DAGH (Figure 5C). With the exception of weak ^1^H_N_ dispersion for F33, all of the residues in the B and C strands have flat relaxation profiles and exhibit neither ^15^N nor ^1^H_N_ dispersion (Figure S8). Perturbations in the hydrogen bonding network are largely restricted to strands E and F, with the largest perturbations (ΔωN > 1.8 ppm) occurring in edge strand F, which is in the strong dimer interface in the TTR tetramer where its C-terminal side chains form part of the hydrophobic pocket that accommodates the F87 ring from the neighboring protomer. The conformational fluctuations observed in the F strand of monomeric F87E, which propagate through the hydrogen bonding network to strand E, likely reflect loss of stabilizing interactions with the F87 aromatic ring and the F strand of the neighboring protomer upon dissociation of the tetramer.

The relaxation dispersion data reveal conformational exchange processes throughout the AB loop and the EF helix (Figure 5D). Within the TTR tetramer, the AB loop forms the weak dimer interface, packing against the GH loop and C-terminal region of the H strand of the neighboring dimer. Not surprisingly, loss of these interactions in the F87E monomer destabilizes the AB loop, which undergoes conformational fluctuations that lead to relaxation dispersion. ^13^C dispersion is observed for the carbonyls of residues 17-22, indicating backbone conformational changes between the ground and excited states. Residues 18, 19, 22 and 23 exhibit ^15^N or ^1^H_N_ relaxation dispersion indicative of changes in the backbone hydrogen bonding network of the AB loop (Figure 5D). The conformational fluctuations in the AB loop appear to be coupled to fluctuations in the EF helix. The side chains of W79, L82, and I84 are in direct contact with V20 and R21 and the Y78 hydroxyl group hydrogen bonds to the Oδ2 carboxyl oxygen of D18, while Oδ1 is within hydrogen bonding distance of the V20 and R21 amide protons. All residues in the EF helix exhibit ^15^N, ^1^H_N_, or ^13^C’ dispersion indicative of an altered conformation and perturbed hydrogen bonding in the excited state (Figure 5D). In particular, upfield shifts of the ^15^N and ^1^H_N_ resonances of Y78 (ΔωN = −2.17ppm, ΔωH = −0.42 ppm) suggest weakening of critical hydrogen bonds that likely destabilizes the helix in the excited state.

## Discussion

Aggregation and amyloidosis of TTR proceed through dissociation of the native tetramer and misfolding of the monomers to form a partially unfolded amyloidogenic intermediate.*^6,42,43^* To elucidate the atomic-resolution structural changes that are responsible for the conversion of this highly stable serum protein into an amyloidogenic species, we have characterized the solution structure and conformational fluctuations of a designed TTR monomer in which the critical F87 side chain, which contributes strongly to tetramer stability,*^44^* is replaced by glutamate. Insights into the ground state structure of the F87E monomer were derived from an analysis of chemical shifts and shift differences relative to the TTR tetramer. Our strategy to characterize the amyloidogenic intermediate focused on determining excited state chemical shifts of F87E using NMR relaxation dispersion experiments, which require a protein construct from which high-quality NMR data can be collected, but which retains the amyloidogenic behavior of the native protein. Relaxation dispersion experiments typically require several days, even with rapid data acquisition techniques, so samples must remain stable over this time period. The F87E mutant fulfils these requirements and yields high-quality spectra and relaxation dispersion data that enable us to better define the kinetic parameters and extend the sequence coverage relative to previous studies of M-TTR.*^14^* We were also able to extend the analysis to the amide proton and carbonyl carbon resonances to derive a more complete picture of the structural changes in the amyloidogenic excited state.

As a starting point, the structure of the monomeric F87E ground state was elucidated by comparing chemical shifts to those of the wild-type TTR tetramer (Figure 3A). We first probed the conformation of the tetramer in solution by comparing backbone φ and ψ dihedral angles predicted from chemical shifts using TALOS-N*^30^* with backbone dihedral angles in a high resolution crystal structure (1.30 Å, 5CN3)*^12^* of wild-type TTR (Figure S4). Differences between solution and crystal dihedral angles are mostly very small, showing that TTR adopts a very similar secondary structure in the crystal and in solution. Only a single set of resonances is observed for the wild-type TTR tetramer in solution, showing that all four protomers adopt the same conformation. In contrast, crystal structures of tetrameric TTR in the P2_1_2_1_2 space group contain two protomers (chains A and B) in the asymmetric unit that differ in the conformation of the FG loop. The structural differences between the A and B chains in the crystal have been attributed to lattice contacts.*^45,46^* In chain A, N98 is in the α_R_region with a positive LJ dihedral angle while in chain B it lies in the β region of the Ramachandran plot. A single conformation is observed for the FG loop of the TTR tetramer in solution, with predicted LJ and ψ angles that are similar to those of chain A in the crystal structure, but which differ greatly from the angles in chain B (Figure S4).

Comparison of the chemical shifts and predicted backbone dihedral angles for the F87E monomer and the WT TTR tetramer (Figure S5A) shows that their secondary structures are very similar. The A, B, C, D, and E β-strands and EF helix are stably folded and of similar length in both the tetramer and F87E monomer. However, dissociation of the tetramer leads to changes in conformation and dynamics for residues in the subunit interfaces. Cross peaks of residues in the N-terminal region of strand F and in the GH hairpin are exchange broadened, implying enhanced conformational fluctuations in this region caused by the F87E mutation and disruption of the strong and weak dimer interfaces. Conformational changes are also observed in the FG loop, where there are large chemical shift differences and the LJ dihedral angle of N98 shifts from positive (α_R_ region) in the tetramer to the polyproline II region (negative LJ) in the monomeric F87E. Differences in chemical shift and order parameters are observed in the AB loop, EF loop, and F β-strand, indicating changes in conformation or dynamics in these regions upon dissociation of the tetramer (Figure 3A, Figure S5).

The F87E monomer adopts two approximately equally populated conformers in solution, which differ in the structure and interactions of the H strand. In state B, the amide proton resonances of A120 and V121 have small chemical shift temperature coefficients, showing that they participate in hydrogen-bonded secondary structure, probably through interactions with strand G. In state A, by contrast, residues 118-123 exhibit near random coil chemical shifts and reduced chemical shift order parameters (Figure S5B), indicating that the C-terminal end of the H strand is dissociated and is highly dynamic. The F87E structure appears to be overall very similar to the 3D solution structure of M-TTR at 500 bar pressure determined from NOE distance restraints.*^15^* In common with state A of F87E TTR, the H β-strand of M-TTR is not stably formed in the high pressure NMR structure, while strands A and G, the entire CBEF β-sheet, and the EF helix adopt native-like structures.

Previous ^15^N relaxation dispersion measurements on M-TTR at physiological pH revealed local structural fluctuations in the DAGH β-sheet that are enhanced at the acidic pH that promotes amyloid formation,*^14^* but neither the timescale of the conformational fluctuations nor the detailed structural changes that accompany formation of the excited state could be ascertained from ^15^N relaxation dispersion data alone. In the present work, we extended this analysis to the monomeric F87E variant, measuring single, double, and zero quantum relaxation dispersion profiles for ^15^N, ^1^H_N_, and ^13^C’ resonances to better define the kinetic parameters and structural changes associated with intrinsic conformational fluctuations of the TTR monomer.

The relaxation dispersion data probe exchange between the ground state and a higher energy conformational state (excited state) of the F87E monomer and provide important insights into the molecular events that lead to local unfolding and aggregation. Based on recent cryo-electron microscopy structures of patient-derived transthyretin fibrils,*^47–51^* it is clear that complete unfolding and structural rearrangement of the dissociated monomer is required for fibril formation (Figure 6). Thus, understanding of the intrinsic structural fluctuations of the TTR monomer and how they are affected by acidic pH, which promotes partial unfolding and aggregation, is of prime importance.

**Figure 6.**
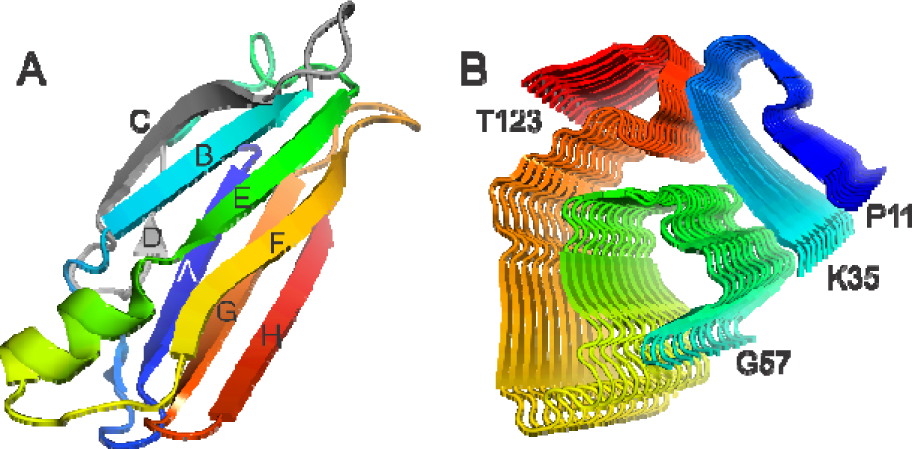
Comparison of the backbone structure of TTR in A. as the folded protomer (subunit from PDB 5CN3)*^12^* and B. the fibrillar form (PDB 6SDZ)*^51^*. The fibril structures (panel B) are shown in rainbow colors, with the N-terminal (P11) blue and the C-terminus red. The colors in panel A correspond to those in panel B; the residues in the C and D strands and the CD loop, which are missing from the fibril structure, are colored gray.

In agreement with relaxation dispersion studies of M-TTR, supported by molecular dynamics simulations, we observe the CBEF β-sheet of the F87E monomer to be more stable than the DAGH sheet, with conformational fluctuations limited to the outer strand F and neighboring E strand.*^14,52^* As in M-TTR,*^14^*conformational fluctuations are observed across the entire hydrogen bonding network of the DAGH sheet of F87E at pH 6.5 (Figure 5A), propagated from the strong dimer interface where loss of the H/H’ interactions of the tetramer leads to structural destabilization. While dissociation of the H strand exposes highly amyloidogenic sequences in strand G (^105^YTIAAL) and strand H (^118^TTAVV),*^51,53^* this appears to be insufficient to drive TTR into the aggregation pathway. The F87E monomer is intrinsically stable and shows no signs of aggregation over a period of several weeks at pH 6.5 - 7, even though there is an approximately 50% population of a state (state A) in which the H strand is dissociated and dynamically disordered. Thus, although the population of state A is increased at acidic pH (Figure S10 and ref.*^40^*), its inherent stability suggests that we must look elsewhere for clues to the local unfolding processes that initiate aggregation.

The ΔωN, ΔωH, and ΔωC’ chemical shift differences derived from fits of the relaxation dispersion data indicate perturbation of the interactions between β strands A and D in the excited state of F87E. The observed ^15^N and ^1^H_N_ chemical shift differences indicate fluctuations, and likely weakening, of critical backbone hydrogen bonding interactions between the A and D strands and weakening or rupture of the C10Sγ - G57 NH hydrogen bond in the excited state. In addition, ^13^C’ dispersion provides evidence of conformational changes in the N-terminal regions of the A and D strands. Upon lowering the pH from 6.7 to 4.4, a pH that strongly promotes aggregation, the intensities of the amide cross peaks of L12, V14, E54, H56, and G57 are strongly attenuated (Table S1, data from ref.*^40^*) indicating enhanced conformational fluctuations or increased population of the excited state in which the interstrand hydrogen bonding network between β-strand D (residues 54-56) and the N-terminal region of strand A is perturbed. Disruption of the interactions between the A and D strand represents a highly plausible mechanism by which aggregation is initiated. Conformational fluctuations that destabilize strand D and lead to its dissociation would expose a strongly amyloidogenic sequence (^12^LMVKV) in strand A. At a later stage, as amyloid formation progresses and strand D becomes fully dissociated, the C-terminal region of strand C would likely be destabilized by disruption of inter-strand packing interactions with strand D side chains. The N-terminal region of strand C appears to retain native-like structure in the pH 4.4 amyloidogenic intermediate*^14,54,55^* but becomes unfolded during fibril formation.*^51^* Approximately 25% of known disease-causing mutations are located in the region encompassing β-strands C and D*^56^* and biochemical, structural, and molecular dynamics studies point to the C/D region as a critical initiation site for partial unfolding and progression down the aggregation pathway.*^57–64^* Residues A36 to H56, spanning β-strands C and D and the BC and CD loops in the native protomer structure, are absent from the structures of TTR fibrils.*^47–51^* Proteolytic cleavage is observed at several sites in the C/D region in patient-derived fibrils, indicating that the entire region is disordered and exposed to proteases in aggregated and fibrillar states of TTR.*^48^*

The AB loop in tetrameric TTR is stabilized by inter-subunit interactions across the weak dimer interface and by intra-protomer interactions with side chains on the EF helix. The EF helix and EF loop act as scaffolds that stabilize both the weak and strong dimer interfaces of the tetramer and are directly implicated in amyloid formation.*^65^* Loss of the inter-subunit interactions, as modeled by the F87E monomer, destabilizes the AB loop and results in conformational fluctuations that are propagated to the EF helix via direct van der Waals contacts and the critical D18 Oδ – Y78 hydroxyl hydrogen bond (Figure 5D). Relaxation dispersion data, acquired at near neutral pH, show that the F87E monomer transiently populates an excited state with a perturbed hydrogen bonding network in both the AB loop and EF helix. Y78 undergoes large upfield chemical shift changes in the excited state (ΔωN = −2.17 ppm, ΔωH = −0.42 ppm), consistent with substantial weakening of hydrogen bonds in the EF helix and destabilization of the monomer structure. The D18-Y78 interaction plays a critical role in stabilization of TTR and mutations at these sites (D18G and Y78F) destabilize both the tetramer and monomer and render the protein highly amyloidogenic.*^65–67^*

The conformational fluctuations in the AB loop and EF helix and loop are enhanced at acidic pH (Table S1), resulting in severe attenuation of the amide cross peak intensities of L17, D18, and V20 in the AB loop and of A81 - I84, the C-cap residues of the EF helix.*^65^* In contrast, the cross peaks of R21 in the AB loop and T75, K76, and Y78 in the EF helix gain intensity under conditions (pH 4.4) where fibril formation is maximal,*^6^* suggesting local unfolding that increases the flexibility of the polypeptide backbone at the N-terminus of the EF helix and in part of the AB loop. Destabilization of the EF helix and loop has also been observed in an X-ray structure of the wild type TTR tetramer at pH 3.5,*^68^* where residues K76 to S85 are disordered and missing from the electron density of chain B. Cryo-electron microscopy structures of patient-derived fibrils show that the EF region becomes remodeled, transitioning from helix in the native TTR tetramer to a pair of short β-strands in the fibril.*^47–51^* Unfolding of the helix exposes side chains that mediate critical packing interactions with a β-strand formed by the F and FG regions of the native protomer. Thus, the conformational fluctuations that destabilize the EF helix in the excited state of the TTR monomer and promote local unfolding at acidic pH would facilitate structural remodeling and progression towards amyloid fibrils.

Finally, conformational fluctuations were observed in the DE loop (residues 59-66) of F87E TTR. The relatively large values of ΔωN and ΔωH for several residues (Figure 5B) indicate substantial remodeling of this loop in the excited state. The conformational malleability of the region is evident in recent cryo-electron microscopy structures of cardiac fibrils derived from patients with the I84S mutation, where residues L58 to G67 adopt different conformations in polymorphic structures and function as a gate that opens or closes a central polar channel, found in all TTR fibril structures determined to date.*^47^* Structural heterogeneity has also been observed in the gate region of patient-derived V30M fibrils.*^48^*

Based on mutagenesis and peptide studies, it has been suggested that β-strands F and H play an active role in TTR aggregation.*^53^* However, the current studies of the F87E variant show that release of the H strand by itself is insufficient to initiate aggregation. Although a peptide derived from strand H spontaneously assembles to form a steric zipper,*^53^* self-assembly was not observed in F87E and additional unfolding events were required to drive aggregation. It is reasonable to suppose that, under more strongly amyloidogenic conditions or seeding by preformed fibrils,*^69^* enhanced conformational fluctuations of strand F, at the edge of the CBEF sheet (Figure 5C), may lead to strand dissociation and aggregation.

In summary, the present work provides important insights into the molecular processes that promote local unfolding of the transthyretin monomer and progression along the pathway to amyloid fibrils. While the monomer is stable at neutral pH, conformational fluctuations of the ground state structure populate an excited state in which the hydrogen bonding networks in the DAGH β-sheet, β-strand F, and the AB loop and EF helix are perturbed. The fluctuations are enhanced under conditions that promote amyloid formation, such as weakly acidic pH, suggesting a highly plausible model for aggregation and fibril formation. In this model, the increased amplitude of fluctuations under conditions that favor amyloid formation leads to dissociation of β-strands D, H, and F from the edges of the DAGH and CBEF β-sheets, exposing strongly amyloidogenic amino acid sequences that drive self-association and entry into the aggregation pathway. Edge strand dissociation would destabilize the β-sheets and facilitate the structural remodeling that is observed in patient-derived fibrils.

## Methods

### Sample preparation

#### Expression and purification of TTR F87E

The F87E mutation was introduced into the coding sequence of TTR in pET 29c using Quikchange mutagenesis (Agilent). For protonated samples, BL21 star (DE3) cells (Thermo-Fisher Scientific) were transformed with TTR F87E/pET 29 and plated on LB plates supplemented with kanamycin. 50 ml starter cultures, inoculated from a single colony, were grown overnight in M9 media at 37°C and used to inoculate large cultures, typically 0.5 L. Cells were grown to OD_600_ of 0.8 and induced with 1 mM IPTG at 25°C for 16 hrs. Cells were harvested by centrifugation and frozen at −20°C. For deuterated samples, a 2 ml LB culture supplemented with 50mg/L of kanamycin was inoculated from a single colony and grown to OD_600_ of 0.5. Cells were adapted to 100% deuterated media by addition of 1 ml of culture to 3 ml of M9 media made with successively higher percentages (50%, 75%, 90% and 98%) of D_2_O. Perdeuterated samples were made using deuterated glucose in the media. For purification, cells were resuspended in 40 ml Tris-buffered saline (TBS) buffer with 1 tablet of Pierce Protease inhibitor (Thermo-Fisher Scientific). Cells were lysed by sonication (30 seconds at 40% power, intervals of 1 second with 1 second pause) followed by centrifugation for 20 minutes at 23,000 g. Impurities were precipitated by addition of 8.5g of ammonium sulfate and gently stirring at 4°C for 20 minutes followed by further centrifugation at 23,000 g. F87E TTR was precipitated by addition of a further 13.5 g of ammonium sulfate and stirring at 4°C for 20 minutes followed by centrifugation for 20 minutes at 23,000 g. For perdeuterated samples, the initial ammonium sulfate precipitation was omitted, and impurities were removed using a Superdex 75 column (GE Life Sciences). Protein pellets were resuspended in 4 ml of TBS buffer with protease inhibitor and further purified by size-exclusion chromatography on a Sephacryl 26/60 column pre-equilibrated with buffer containing10mM potassium phosphate, 100mM potassium chloride, 1 mM EDTA, pH 7. Fractions were analyzed by SDS-PAGE and protein rich fractions were pooled and diluted by addition of 2 volumes of 20mM Tris-HCl, pH 8 for ion exchange chromatography. Buffers were (A) 25 mM Tris-HCL pH 8, 1mM EDTA and (B) 25mM Tris-HCl pH 8, 100mM NaCl, 1mM EDTA. Pooled fractions were loaded onto a 5 ml Q-HP column (GE Life Sciences) equilibrated in buffer A and washed with 5% buffer B. TTR was eluted with an initial gradient from 5% to 35% buffer B, which was maintained until the protein had eluted, around 3 column volumes. For NMR spectroscopy, protein-rich fractions were exchanged into buffer containing 50 mM Bis-Tris/MES and 100 mM NaCl, pH 6.5. Except where noted, all samples were concentrated to 300 μM TTR in Vivaspin 20 concentrators (Sartorius) and added to 5 mm susceptibility-matched Shigemi tubes with 5% D_2_O.

#### Aggregation assays

Stock solutions of TTR variants were filtered through a 0.22 µm syringe filter and diluted to a final concentration of 0.2 mg/ml in either 10 mM sodium phosphate buffer at pH 7, or 10 mM sodium acetate buffer at pH 4.3. Both buffers contained 100 mM KCl and 1 mM EDTA, and samples were prepared with and without 1 µM thioflavin T (Sigma). Four separate samples (100 µl) of each TTR solution were incubated in a black, clear-bottom 96-well plate (Corning, Tewksbury, MA, item #3631) at 37 °C in a Spectramax Gemini EM plate reader (Molecular Devices, Sunnyvale, CA). Absorbance was measured at 330 nm, following which the plate was sealed and fluorescence measurements started. Thioflavin T fluorescence (λ_ex_= 440 nm, λ_em_= 480 nm) was measured every 10 minutes. The plate was shaken for 5 s before each reading but otherwise the reactions were not agitated. After 24 h, the absorbance at 330 nm was measured again and the difference recorded. Samples with and without thioflavin T had similar absorbance changes, confirming that the dye did not affect bulk aggregation.

#### NMR Spectroscopy

All NMR data were acquired at 298K on Bruker Avance spectrometers operating at 500, 700, 800 and 900 MHz. The 500 MHz and 800 MHz spectrometers used for relaxation dispersion measurements were equipped with cryoprobes, and the 900 MHz spectrometer was equipped with a triple-axis gradient probe, facilitating direct calibration of gradients for diffusion experiments. All data were collected using XwinNMR and Topspin, processed using NMRPipe*^70^* and analyzed using NMRFam Sparky.*^71^*

#### Resonance assignments

Assignments for ^15^N, ^13^C-labeled F87E TTR were performed using standard triple resonance experiments acquired on a Bruker Avance 700 MHz spectrometer using random non-uniform sampling. The following triple resonance data sets were recorded with non-uniform sampling in the ^15^N and ^13^C dimensions: HNCO, 40% sampling; HNCA, 15% sampling; HNCOCA, 15% sampling; HNCACB, 15% sampling. These spectra were reconstructed using multi-dimensional decomposition (MDD) and co-processed using the mddNMR software package.*^27,28^* Amide cross peak assignments could not be made for contiguous residues between 86 - 92, and between 109 - 116; the cross peaks of these residues appear to be weak and broad and no connectivities could be observed in the triple resonance spectra. Resonance assignments for F87E TTR have been deposited in the Biological Magnetic Resonance Bank (BMRB) with accession code 51171.

Assignments for the wild-type TTR tetramer at pH 6.5 were transferred from those at pH 7 (BMRB 27514). Assignments were checked using carbonyl carbon chemical shifts obtained from standard HNCO experiments.

#### 19F NMR

C10S-S85C-F87E TTR was expressed, purified, and labeled with BTFA using previously described methods.*^72^*

#### Diffusion experiments

Pulsed-field gradient spin-echo diffusion experiments*^24^* were conducted on a Bruker Avance spectrometer operating at 900 MHz equipped with a triple-axis gradient probe. The x-gradient strength (in Gauss) was calibrated from the width of the echo in a spin echo experiment with 10% gradient strength. The diffusion constant is determined by plotting ln(Intensity) against G^2^/cm^2^ and calculating the slope using the Stejskal–Tanner equation.*^23^*

#### NMR Relaxation Dispersion

Relaxation compensated constant time Carr-Purcell-Meiboom-Gill (CPMG) relaxation dispersion experiments*^73,74^* for ^1^H_N_ and ^15^N_H_ resonances were carried out on ^2^H-^15^N samples of TTR F87E with ∼70% deuteration at pH 6.5 and 298K. Data were acquired on 500 and 800 MHz spectrometers using 60% non-uniform sampling in the indirect dimension and a sampling scheme derived using the protocol from Aoto et al.*^75^* with 16 scans per sampling point. In all experiments, heat-compensation was applied after data acquisition and before the inter-scan relaxation delay. ^1^H/^15^N TROSY-based double- and zero-quantum relaxation-compensated constant-time CPMG relaxation dispersion experiments*^35^* were carried out on 200 μM samples of perdeuterated ^2^H-^15^N TTR F87E, using the same sampling scheme as the single quantum experiments. Carbonyl ^13^C’ relaxation dispersion was measured for uniformly ^13^C, ^15^N labeled samples of F87E using the method of Lundstrom et al.*^39^*

Spectra were reconstructed from the non-uniformly sampled data using mddNMR.*^27^* Peak intensities were extracted by fitting peak shapes using FuDA (http://pound.med.utoronto.ca). The ^1^H_N_ and ^15^N_H_ single, double and zero quantum relaxation dispersion profiles were fitted simultaneously to the Carver-Richards equation using the in-house program GLOVE.*^37^* Initially, a set of 15 well-defined curves with minimal random scatter were selected and fitted as a cluster to define global exchange parameters *k_ex_* and *p_B_*. The dispersion profiles for the remaining residues were then fitted using these global parameters. Uncertainties in the extracted chemical shifts (Δω) were estimated using 200 cycles of Monte-Carlo simulations. For a small subset of residues with a large amount of random scatter and high R_2_^0^ for ^1^H_N_ coherences at 800 MHz, only the 500 MHz dispersion profiles plus the ^15^N SQ dispersion profile at 800 MHz were fitted. For two residues (93 and 96), the double- and zero-quantum dispersion profiles could not be fitted globally with the single quantum profiles, so these were excluded from the global fit.

## Supporting information

The supporting information contains Supplementary Tables S1 and S2 and Supplementary Figures S1 – S10.

## Funding

This work was supported by National Institutes of Health Grants DK34909 and DK124211 (PEW) and GM131693 (HJD) and the Skaggs Institute for Chemical Biology. X.S. acknowledges past fellowship support from American Heart Association grants #17POST32810003 and #20POST35050060.

## Acknowledgements

We thank Marga Gairi for assistance with mddNMR, Yvonne Eisele for the TTR used in aggregation assays, Euvel Manlapaz for technical assistance, and Jeffery Kelly for valuable discussions.

## Notes

https://bmrb.io/data_library/held.shtml#51171

### Competing Interest Statement

The authors have declared no competing interest.

