## Supplementary Material for "Characterization of an amyloidogenic intermediate of transthyretin by NMR relaxation dispersion"

[4]Present address: Department of Medicine, Boston University School of Medicine, Boston, Massachusetts 02118.

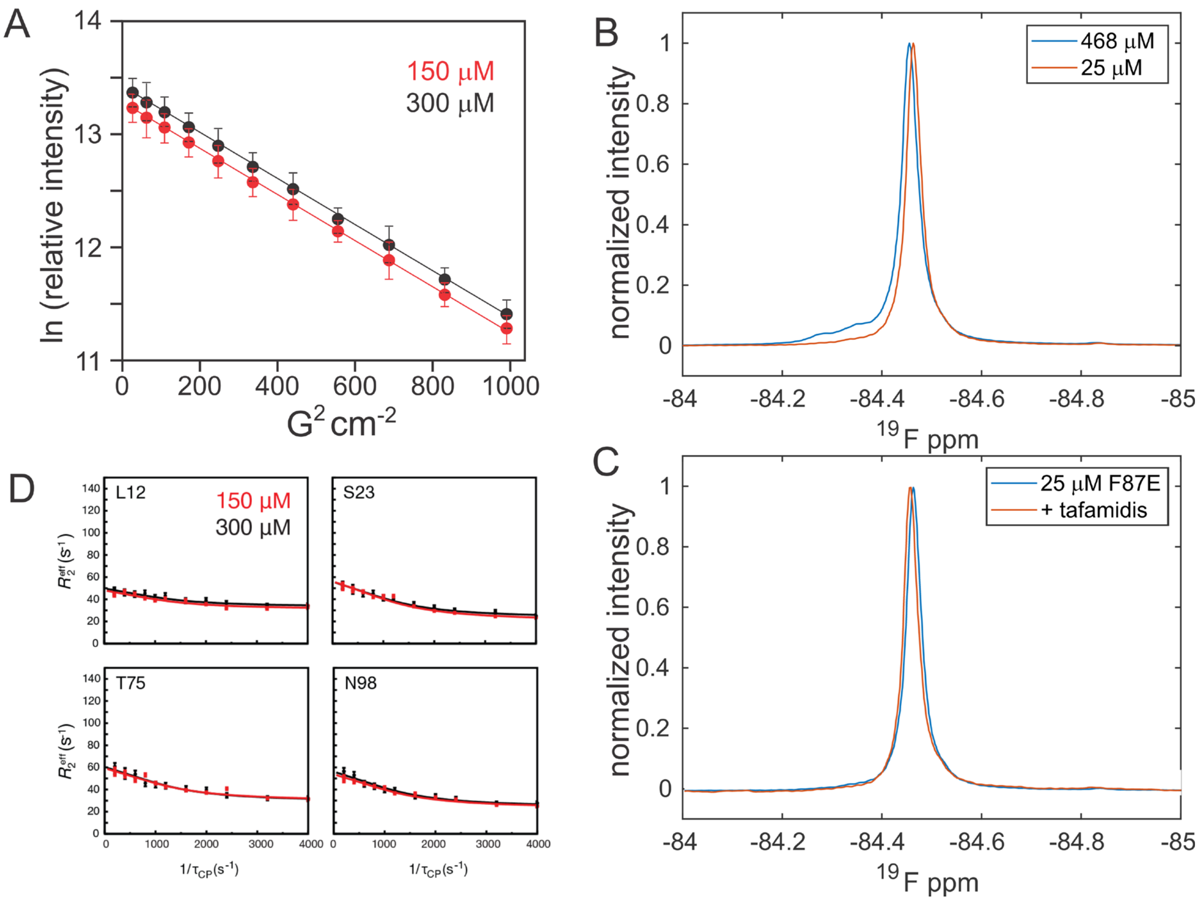

Figure S1. Concentration dependence of F87E mutant. A. Pulsed field gradient spin echo experiment on F87E at 150 μM (red) and 300 μM (black). The solid lines are linear fits. B. $^{19}$F NMR spectra of BTFA-labeled C10S-S85C F87E TTR at 25 μM (red) and 468 μM (blue). C. $^{19}$F NMR spectra of BTFA-labeled C10S-S85C F87E TTR at 25 μM (blue) and with the addition of a 2-fold molar excess of tafamidis (red). D. $^{1}H_{N}$ single quantum relaxation dispersion profiles for representative residues at 150 μM (red) and 300 μM (black).

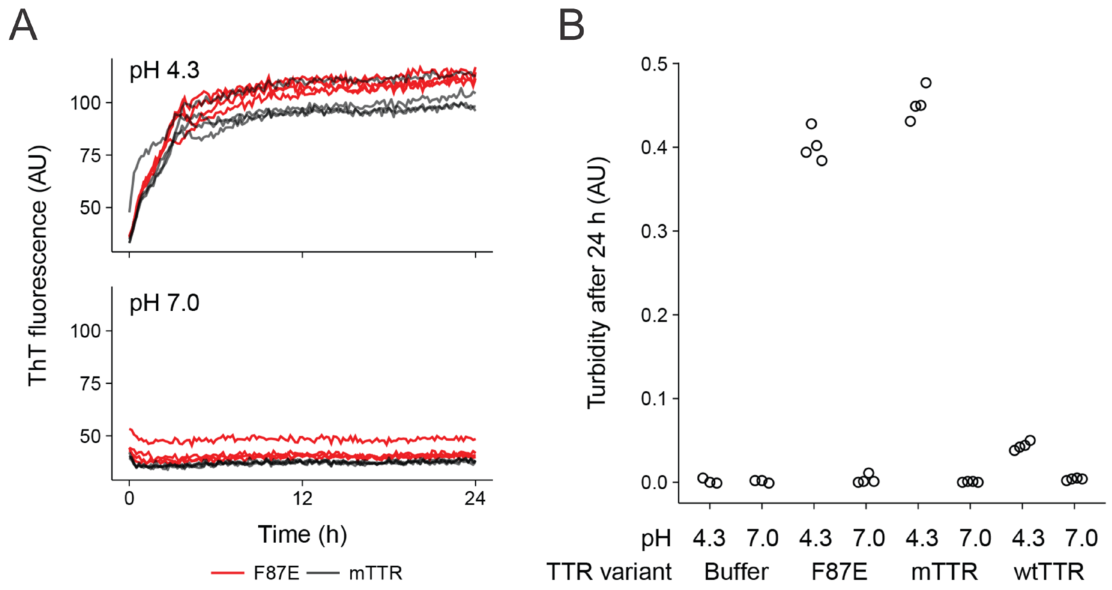

Figure S2. A. Aggregation kinetics for F87E and M-TTR at pH 4.3 and pH 7.0 monitored by thioflavin T fluorescence. Kinetic traces are shown for three replicates for each protein. B. Endpoint stagnant turbidity at 24 hrs for F87E, M-TTR and wild type TTR at pH 4.3 and pH 7.0.

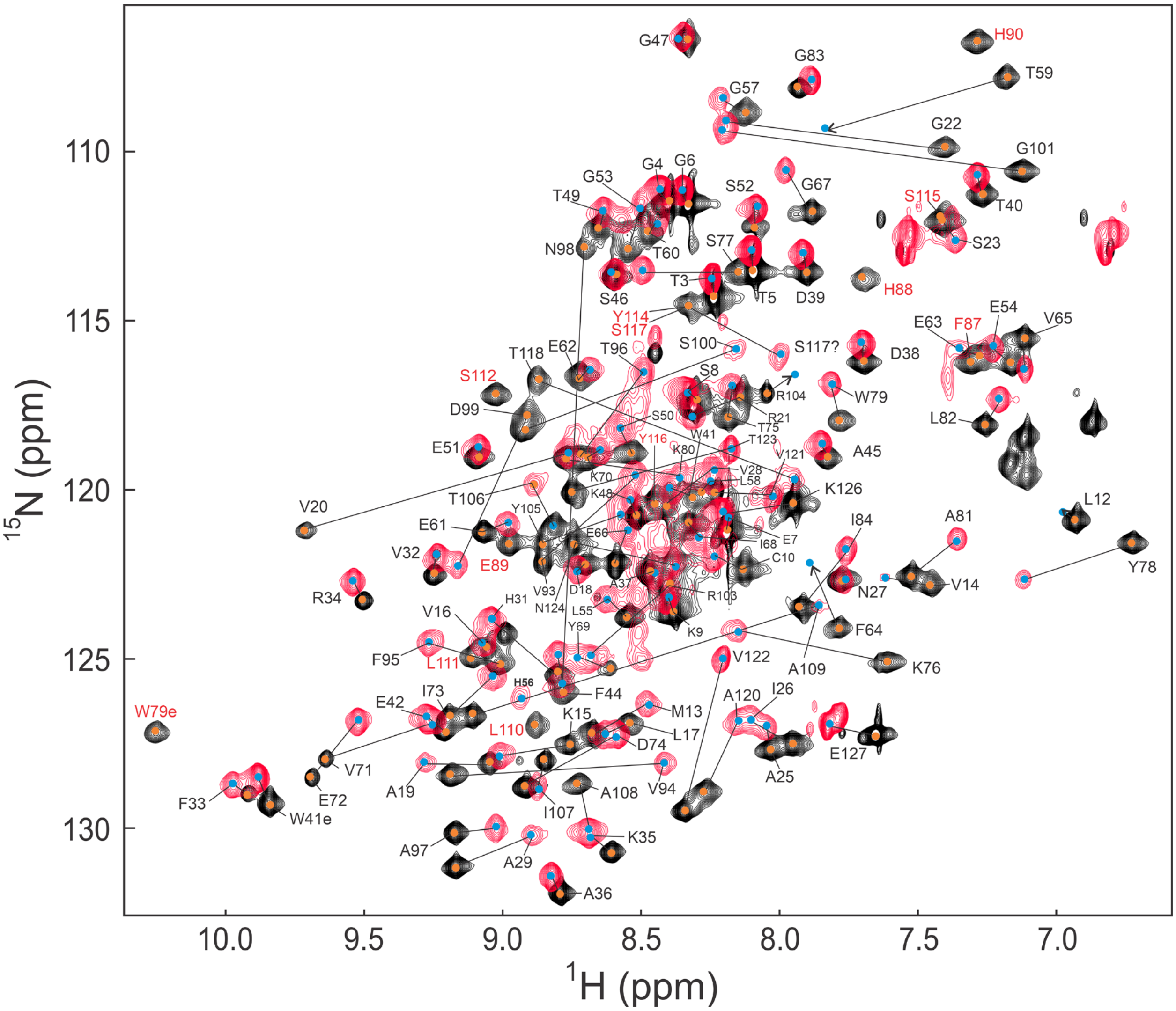

Figure S3. Overlay of the $^{1}$H,$^{15}$N TROSY-HSQC spectrum of WT TTR tetramer (black) and of the $^{1}$H,$^{15}$N HSQC spectrum of F87E TTR (red) at pH 6.5 and 298K. Orange dots denote the location of the TTR tetramer cross peaks, blue dots denote the location of the F87E cross peaks. Corresponding cross peaks are joined by a line. The positions of weak cross peaks in the F87E spectrum, not visible at this contour level, are shown as blue dots and arrows. Cross peaks in the TTR tetramer spectrum that do not appear in the F87E spectrum are labeled in red. Only one cross peak (H56) appears in the F87E spectrum and not in the TTR tetramer spectrum.

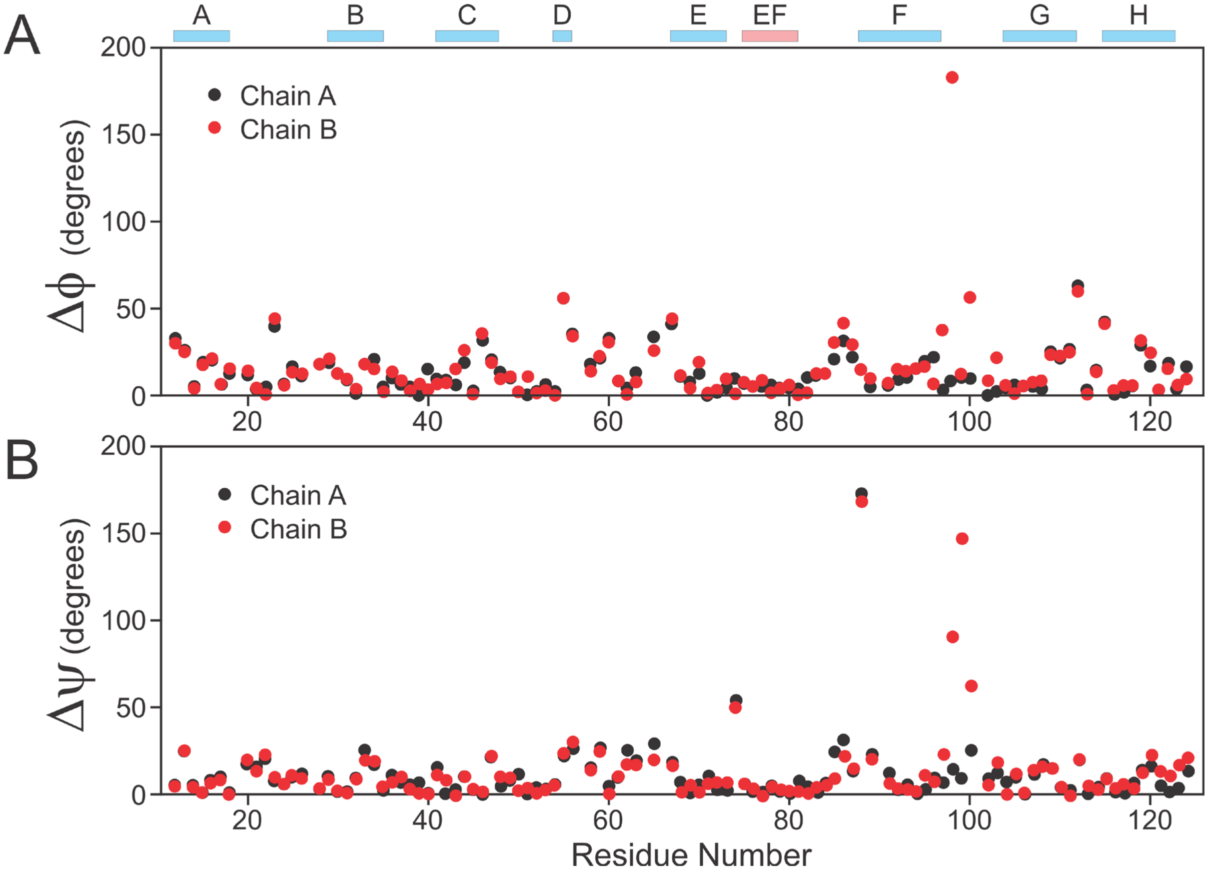

Figure S4. Per-residue difference in the dihedral angles ϕ (A) and ψ (B) between those predicted from the NMR chemical shifts using TALOS-N[1] and those observed in the X-ray crystal structure (PDB 5CN3), chain A (black) or chain B (red) of the asymmetric unit. The colored rectangles at the top of the figure indicate the locations of secondary structure elements in the X-ray crystal structure, β-strands A-H (blue) and the EF-helix (pink).

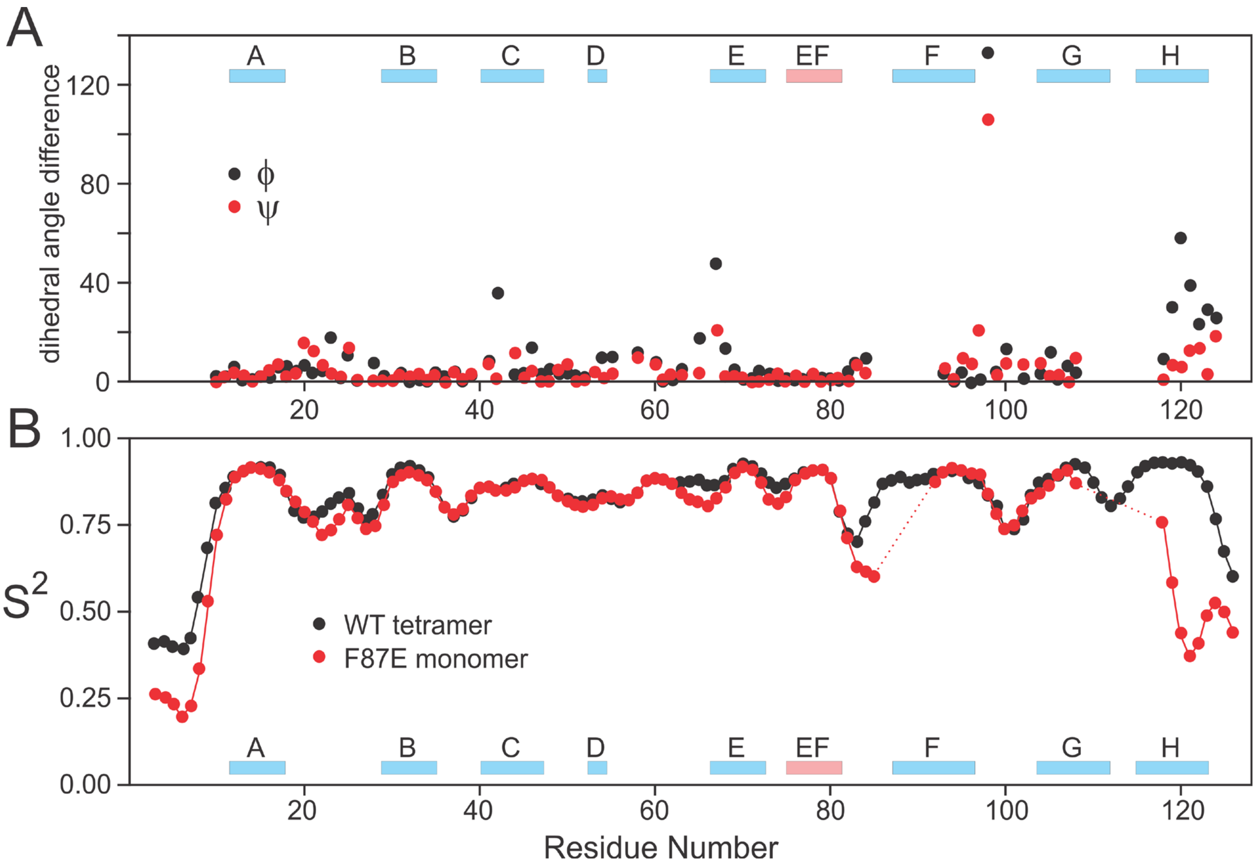

Figure S5. A. Absolute value of the difference in the dihedral angles ϕ (black) and ψ (red), predicted from chemical shifts using TALOS-N[1], between the WT tetramer and F87E. B. RCI order parameters ($S^2$)[2] calculated from the NMR chemical shifts using TALOS-N, reveal increased backbone flexibility in several regions of the F87E monomer (red) relative to the tetramer (black). The colored rectangles at the top and bottom of the figure indicate the locations of secondary structure elements in the X-ray crystal structure, β-strands A-H (blue) and the EF-helix (pink).

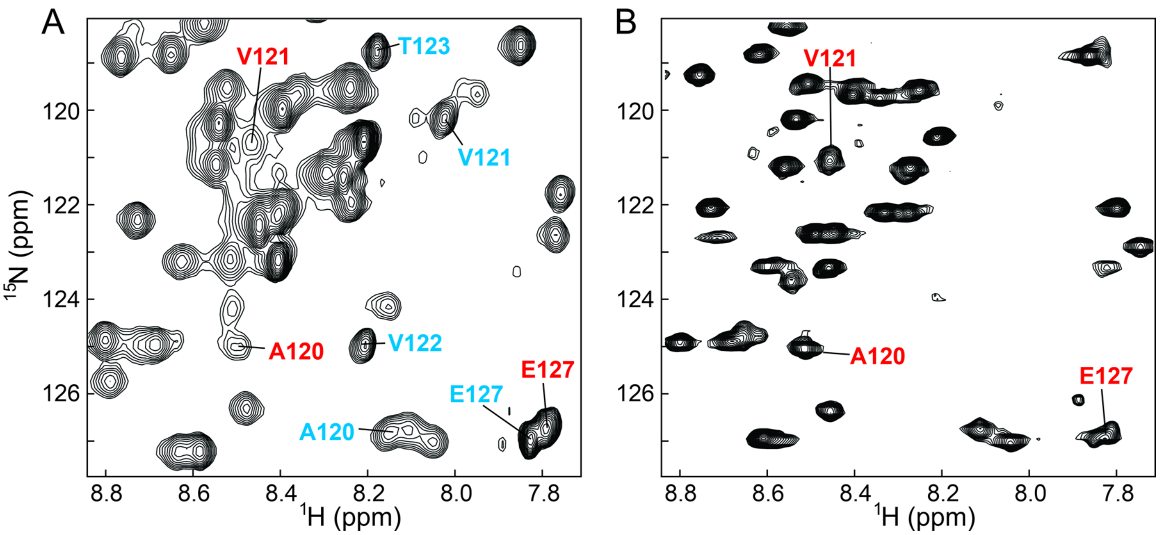

Figure S6. A. Region of the HSQC spectrum of the F87E showing amide cross peaks arising from the two conformations of the H strand. Assignments of residues in state A, with near random coil $^{1}H_{N}$ and $^{15}N$ chemical shifts, are labeled in blue and the cross peaks of A120, V121, and E127 in state B are labeled in red. B. Corresponding region of the HSQC spectrum of M-TTR, with cross peaks of A120, V121, and E127 labeled.

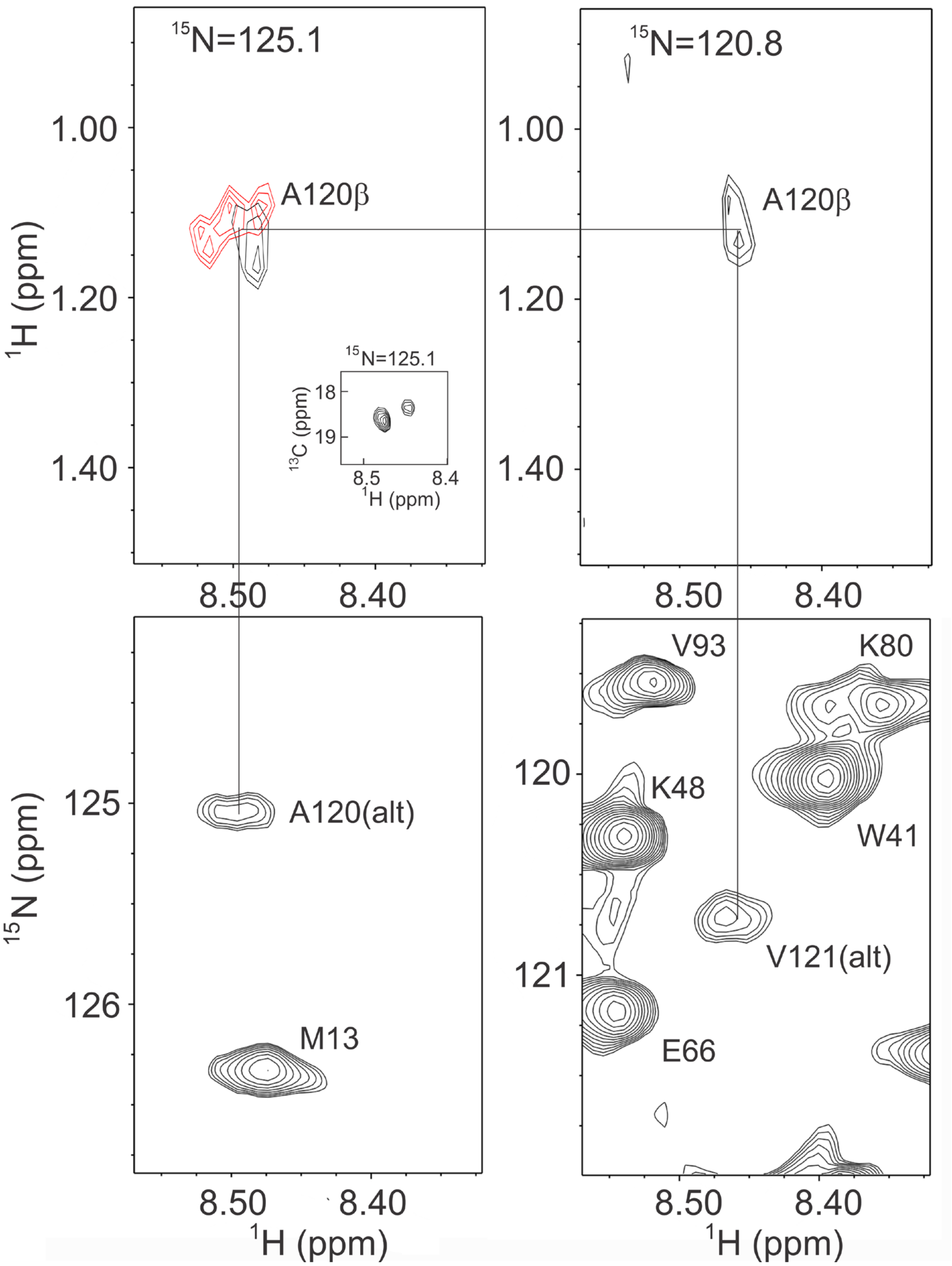

Figure S7. Local regions of planes from a 3D $^{15}$N-edited NOESY-HSQC spectrum (900 MHz, 100 ms mixing time; black cross peaks) showing NOEs from the A120 ($^{15}$N = 125.1 ppm) and V121 ($^{15}$N = 120.8) $^{1}H_{N}$ resonances to a methyl resonance at 1.13 ppm. Both amides also have NOEs to an Hα at 4.86 ppm. A plane ($^{15}$N = 125.1 ppm) from a $^{15}$N-edited TOCSY-HSQC spectrum showing a scalar connectivity between the A120 $^{1}H_{N}$ and the methyl is overlaid in red. Inset: Region of the $^{15}$N = 125.1 ppm plane from a 3D HNCB spectrum showing a cross peak between the A120 amide and a Cβ resonance at 18.7 ppm.

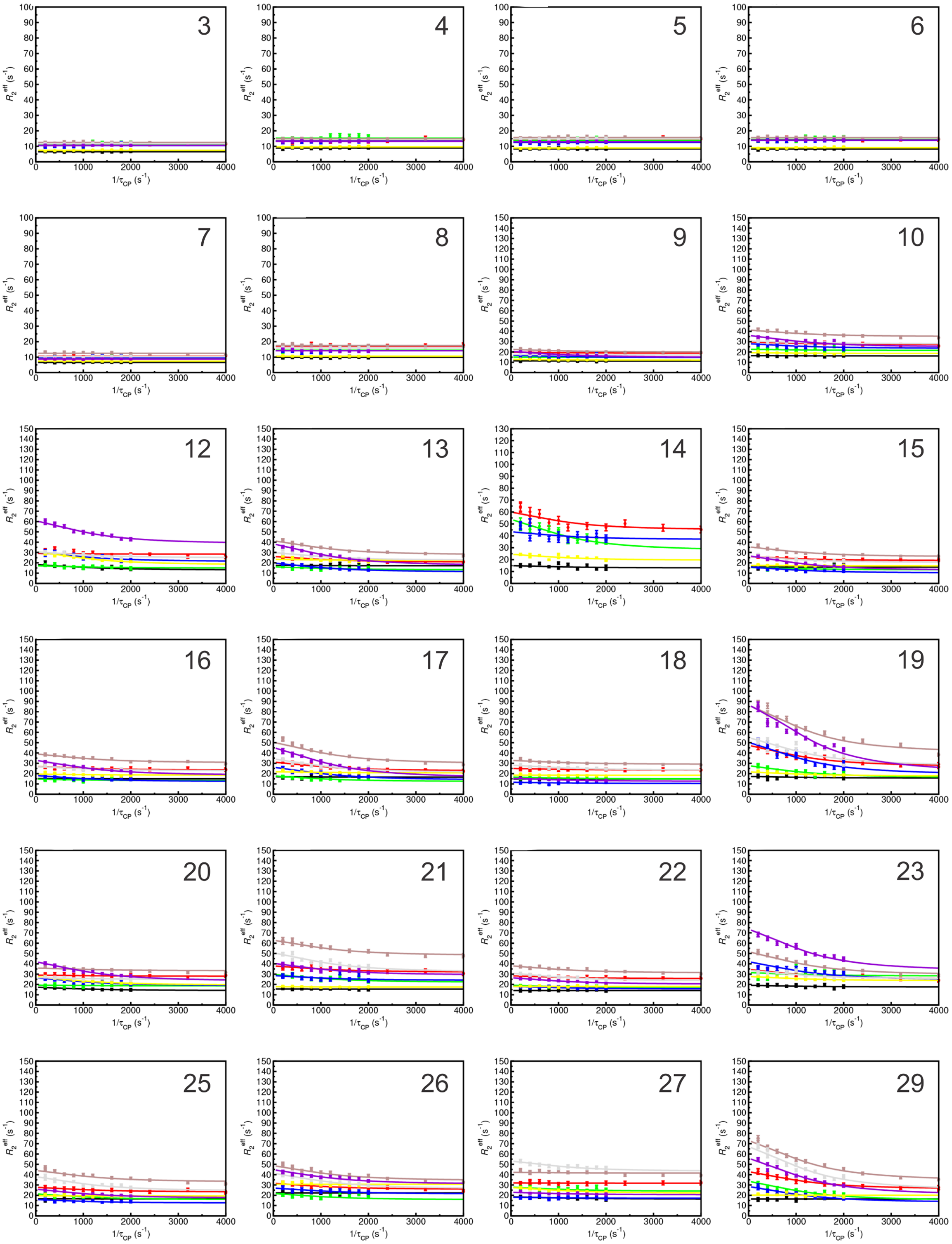

3
4
5
6
7
8
9
10
25
26
27
29
$R_2^{eff}$ (s$^{-1}$)
1/$\tau_{CP}$ (s$^{-1}$)

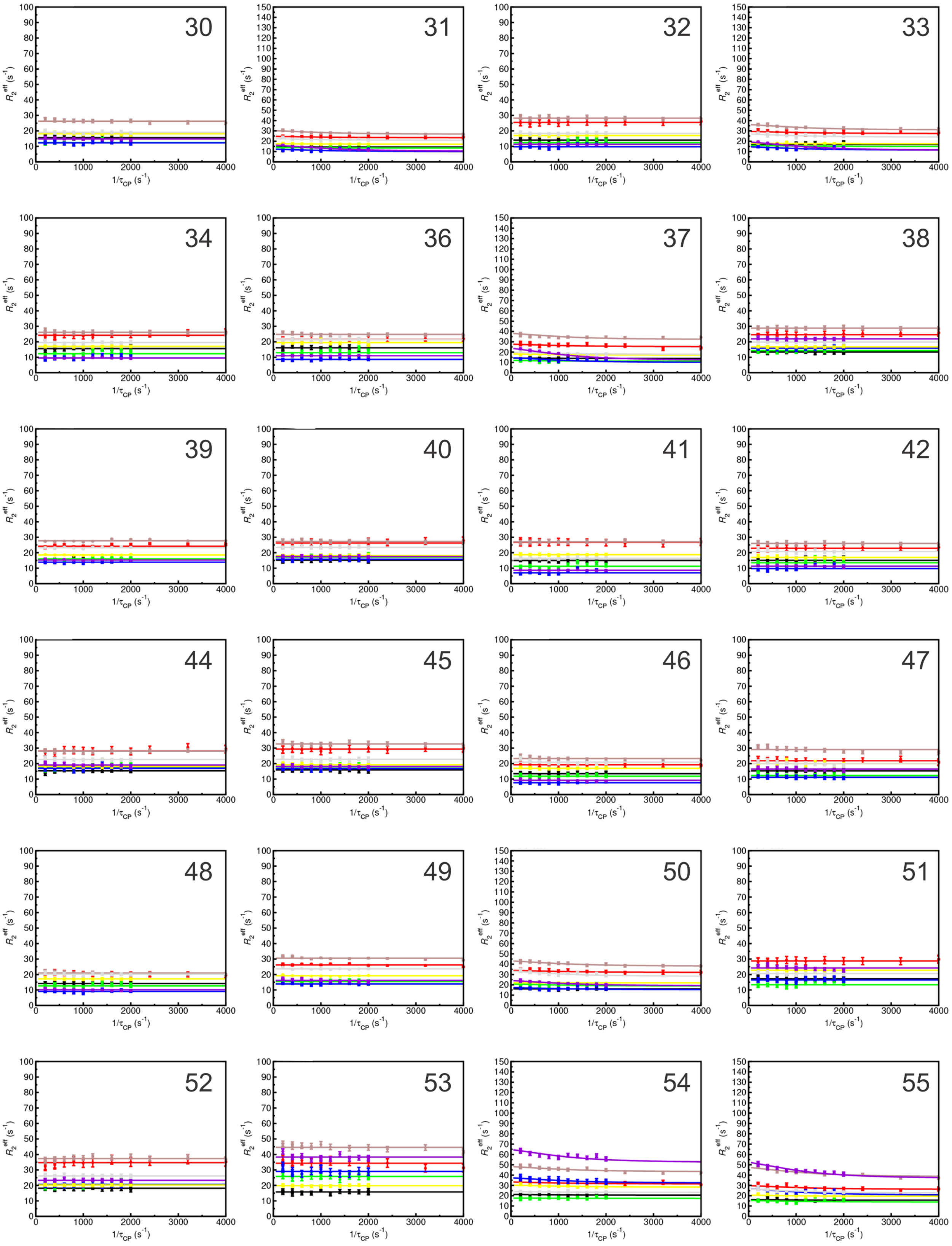
30
31
32
33
34
36
37
38
39
40
41
42
$R_2^{eff}$ ($s^{-1}$)
$1/\tau_{CP}$ ($s^{-1}$)

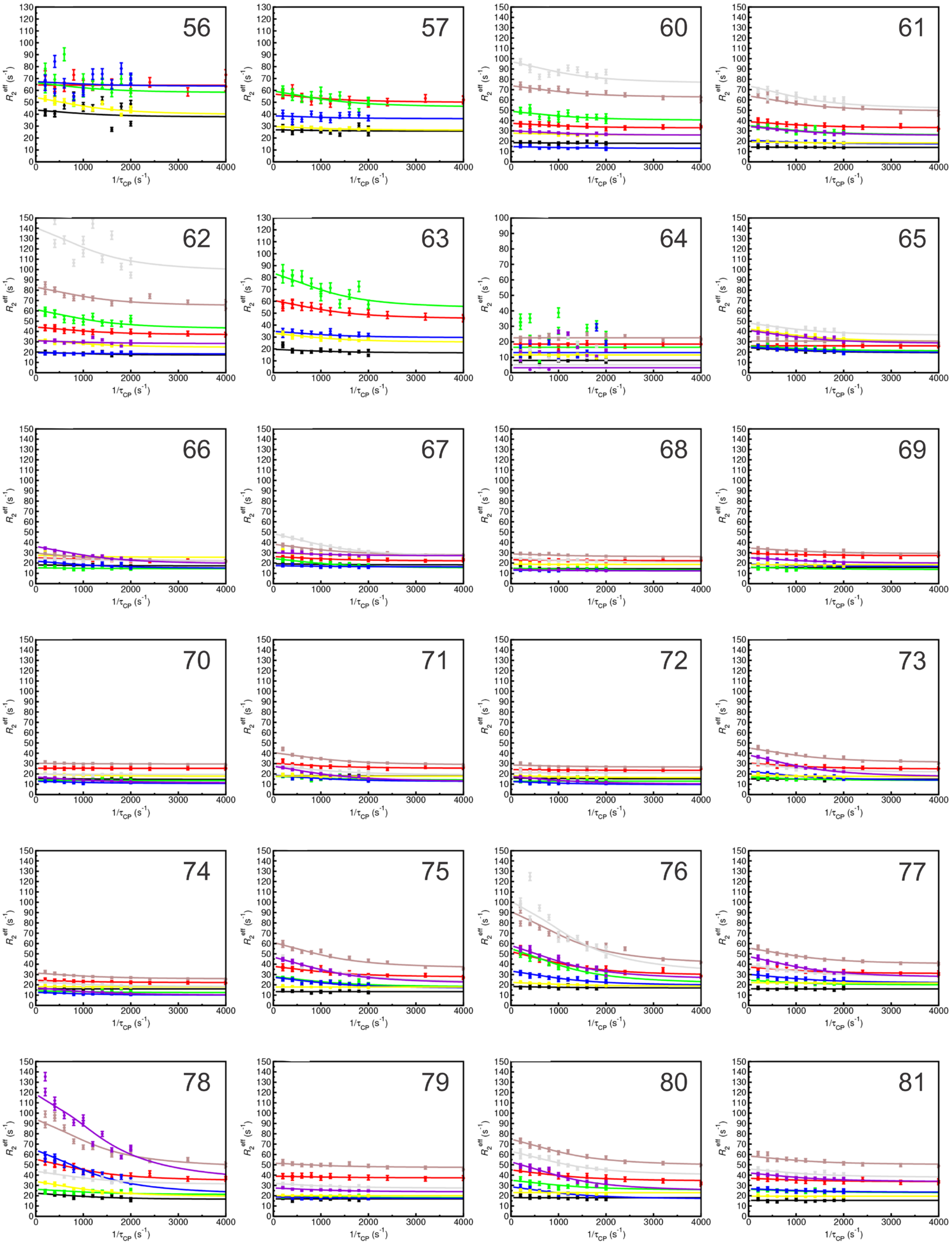

56
57
$R_2^{eff}$ (s$^{-1}$)
1/$\tau_{CP}$ (s$^{-1}$)

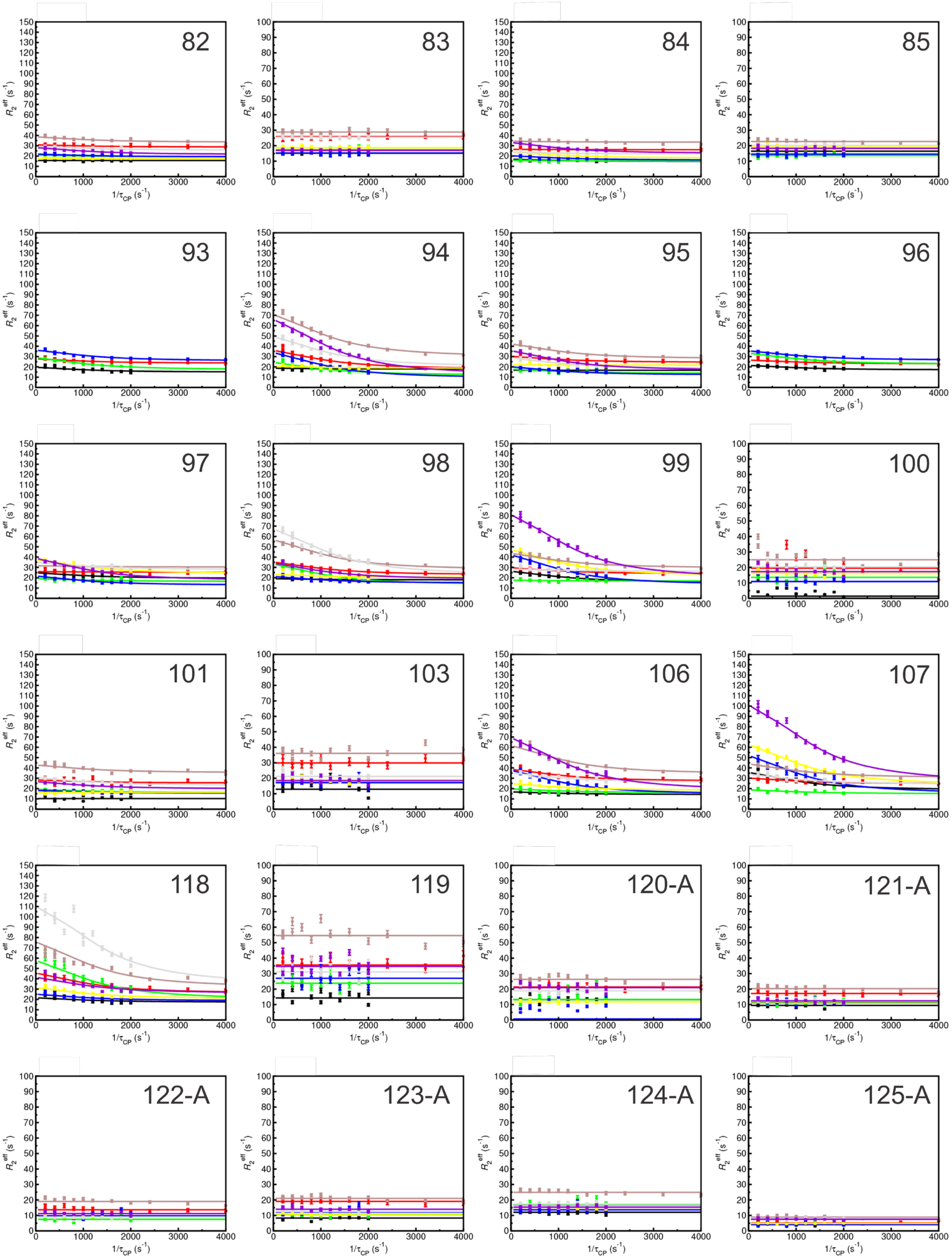
82
83
84
85
93
94
95
96
97
98
99
100
101
103
106
107
118
119
120-A
121-A
122-A
123-A
124-A
125-A
$R_2^{eff}$ ($s^{-1}$)
$1/\tau_{CP}$ ($s^{-1}$)

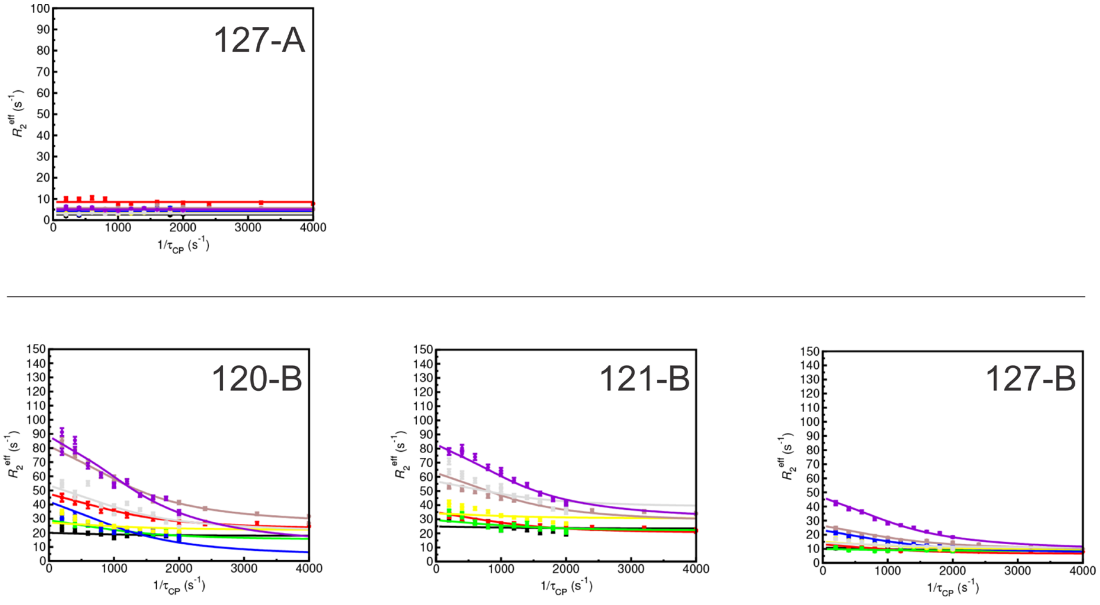

Figure S8. Relaxation dispersion profiles for all assignable and non-overlapped resonances in the spectra of F87E at 800 MHz and 500 MHz. Dispersion profiles for residues 120-127 in state A, where the C-terminal region is dynamically disordered, are shown in the upper panel, above the line. Profiles for residues 120, 121, and 123 in the closed state B are shown in the lower panel. The dispersion profiles are color coded as: red, 500 MHz $^{1}$H single-quantum (SQ); brown, 800 MHz $^{1}$H SQ; black, 500 MHz $^{15}$N SQ; yellow, 800 MHz $^{15}$N SQ; green, 500 MHz $^{1}$H - $^{15}$N zero-quantum (ZQ); gray: 800 MHz $^{1}$H - $^{15}$N ZQ; blue, 500 MHz $^{1}$H - $^{15}$N double-quantum (DQ); purple, 800 MHz $^{1}$H - $^{15}$N DQ. The solid lines represent fits to the Carver-Richards equation[3] with fixed $k_{ex}$ (= 3800 s$^{-1}$) and $p_B$ (= 0.054) derived from fitting the $^{15}$N and $^{1}$H$_N$ SQ, ZQ, and DQ dispersion data for a cluster of 15 residues with well-defined dispersion curves and minimal scatter in the data points.

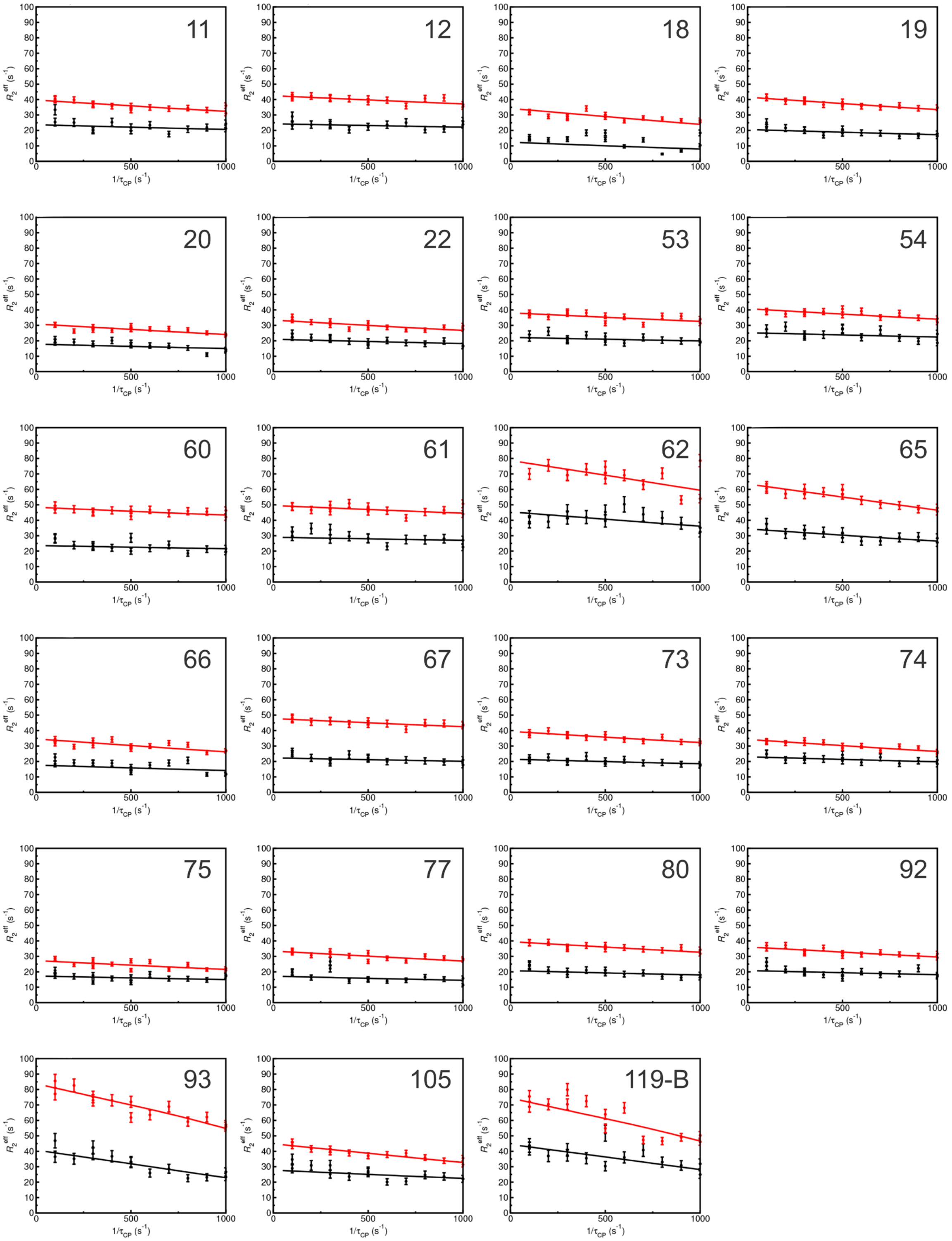

Figure S9. $^{13}$CO relaxation dispersion profiles for F87E recorded at 800 MHz (red) and 500 MHz (black). The solid lines represent fits to the Carver-Richards equation[3] with fixed $k_{ex}$ (= 3800 s$^{-1}$) and $p_B$ (= 0.054) derived from fitting the $^{15}$N and $^{1}H_N$ dispersion data. Residue numbers refer to the carbonyl that exhibits dispersion, detected via the amide of residue i+1 in the peptide bond. Data for residues that do not exhibit dispersion, i.e., have flat dispersion profiles, are not shown.

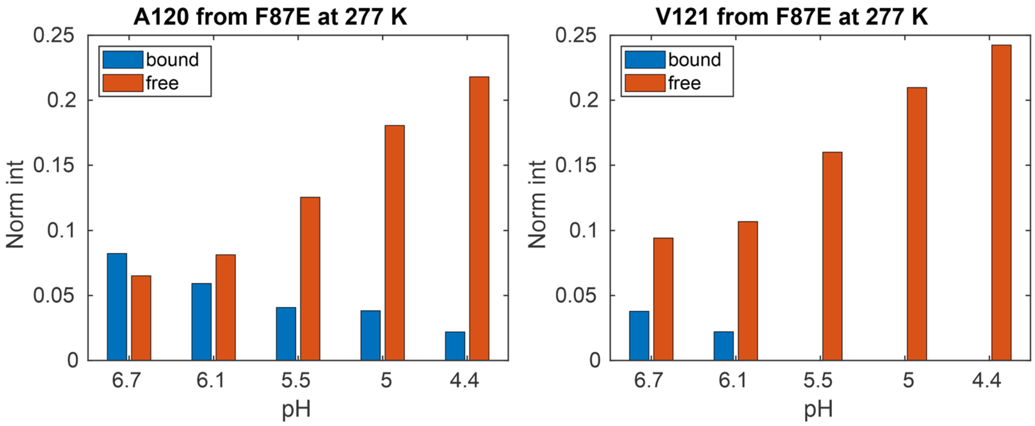

Figure S10. Changes in intensity of A120 and V121 cross peaks in states A (red) and B (blue) of F87E as the pH is reduced from 6.7 to 4.4. The cross peak intensities were measured as peak height and normalized to the average peak height of residues T3, G4, andT5 in the disordered N-terminus.

Table S1. Values of ΔωH and ΔωN from fits of single, zero, and double quantum relaxation dispersion profiles for F87E. Uncertainties were estimated from 200 Monte Carlo simulations. Signs were determined, where possible, from exchange induced chemical shift differences between HSQC and HMQC spectra.

| Residue | ΔωH (ppm) | ΔωH error (ppm) | ΔωN (ppm) | ΔωN error (ppm) | HSQC-HMQC [a] (Hz) | Intensity ratio [b] pH 4.4/pH 6.7 |
|---|---|---|---|---|---|---|
| 9 | 0.09 | 0.007 | 0.44 | 0.07 | | 0.52 |
| ***10*** [c] | ***0.14*** | 0.006 | ***0.40*** | 0.07 | 0.3 | 0.83 |
| 12 | 0.10 | 0.011 | 1.85 | 0.07 | | 0.37 |
| ***13*** | ***-0.19*** | 0.008 | ***0.75*** | 0.08 | 0.6 | 0.52 |
| 14 | 0.34 | 0.020 | 0.75 | 0.09 | | 0.39 |
| 15 | 0.17 | 0.004 | 0.39 | 0.05 | | 0.83 |
| ***16*** | ***-0.16*** | 0.005 | ***-0.57*** | 0.06 | -0.6 | 0.48 |
| 17 | 0.26 | 0.004 | 0.56 | 0.04 | | 0.28 |
| 18 | 0.11 | 0.006 | 0.12 | 0.07 | | 0.32 |
| ***19*** | ***0.42*** | 0.004 | ***1.04*** | 0.04 | 3.0 | 0.92 |
| 20 | 0.09 | 0.007 | 1.55 | 0.06 | | * |
| 21 | 0.22 | 0.006 | 0.25 | 0.07 | | 1.85 |
| 22 | 0.14 | 0.006 | 0.05 | 0.04 | | 0.49 |
| ***23*** | ***0.27*** | 0.005 | ***1.08*** | 0.06 | 1.2 | 0.53 |
| 25 | 0.19 | 0.004 | 0.25 | 0.05 | | 0.59 |
| 26 | 0.21 | 0.005 | 0.06 | 0.05 | | * |
| 27 | 0.05 | 0.010 | 1.28 | 0.08 | | 0.07 |
| 29 | 0.37 | 0.005 | 0.11 | 0.06 | | < 0.3 |
| 31 | 0.10 | 0.008 | 0.36 | 0.08 | | 0.52 |
| ***33*** | ***-0.13*** | 0.006 | ***-0.30*** | 0.06 | -0.5 | 0.69 |
| 37 | 0.14 | 0.006 | 0.58 | 0.06 | | * |
| 50 | 0.12 | 0.006 | 0.06 | 0.04 | | * |
| ***54*** | ***0.17*** | 0.006 | ***0.55*** | 0.07 | 0.8 | 0.24 |
| 55 | 0.12 | 0.011 | 0.76 | 0.10 | | * |
| 56 | 0.06 | 0.043 | 2.12 | 0.12 | | 0.32 |
| ***57*** | ***0.29*** | 0.008 | ***1.08*** | 0.09 | 0.5 | < 0.3 |
| ***60*** | ***-0.21*** | 0.010 | ***0.99*** | 0.09 | 1.7 | * |
| ***61*** | ***0.22*** | 0.006 | ***-0.57*** | 0.08 | -0.9 | 0.16 |
| 62 | 0.24 | 0.010 | 1.50 | 0.08 | | 0.20 |
| 63 | 0.36 | 0.024 | 1.51 | 0.10 | | 0.40 |
| 65 | 0.01 | 0.004 | 1.98 | 0.06 | | * |
| 66 | 0.16 | 0.005 | 0.75 | 0.06 | | 0.30 |
| ***67*** | ***-0.18*** | 0.006 | ***-0.90*** | 0.08 | -0.7 | 0.41 |
| 68 | 0.09 | 0.006 | 0.23 | 0.07 | | 0.72 |
| 69 | 0.13 | 0.005 | 0.05 | 0.04 | | * |
| 70 | 0.04 | 0.008 | 0.88 | 0.08 | | * |
| 71 | 0.19 | 0.004 | 0.24 | 0.04 | | * |
| 72 | 0.08 | 0.008 | 0.73 | 0.08 | | 0.20 |
| 73 | 0.21 | 0.004 | 0.55 | 0.05 | | 1.52 |
| 74 | 0.13 | 0.005 | 0.16 | 0.05 | | 0.97 |
| 75 | 0.29 | 0.004 | 0.05 | 0.04 | | 4.83 |

| | | | | | | |
|---|---|---|---|---|---|---|
| 76 | 0.45 | 0.006 | 1.12 | 0.06 | | 3.76 |
| ***77*** | ***-0.22*** | 0.006 | ***-0.35*** | 0.06 | -0.4 | 0.48 |
| ***78*** | ***-0.42*** | 0.007 | ***-2.17*** | 0.06 | -1.9 | 1.94 |
| 79 | 0.11 | 0.011 | 0.08 | 0.09 | | 0.61 |
| 80 | 0.29 | 0.005 | 0.22 | 0.07 | | * |
| 81 | 0.16 | 0.008 | 0.07 | 0.06 | | < 0.2 |
| 82 | 0.13 | 0.007 | 0.15 | 0.07 | | 0.12 |
| 84 | 0.04 | 0.007 | 1.32 | 0.06 | | 0.27 |
| ***93*** | ***0.18*** | 0.008 | ***1.88*** | 0.07 | 0.4 | * |
| ***94*** | ***0.39*** | 0.004 | ***0.69*** | 0.05 | 1.5 | 0.91 |
| 95 | 0.21 | 0.004 | 0.37 | 0.05 | | 0.34 |
| 96 | 0.17 | 0.007 | 1.80 | 0.09 | | 0.39 |
| 97 | 0.06 | 0.006 | 2.01 | 0.05 | | 0.61 |
| ***98*** | ***0.31*** | 0.004 | ***-0.90*** | 0.05 | -1.7 | 0.56 |
| ***99*** | ***-0.21*** | 0.006 | ***-2.86*** | 0.06 | -1.4 | * |
| 101 | 0.15 | 0.005 | 0.05 | 0.04 | | * |
| ***106*** | ***0.30*** | 0.005 | ***1.38*** | 0.05 | 0.3 | 0.66 |
| 107 | 0.20 | 0.005 | 3.70 | 0.06 | | 1.32 |
| ***118*** | ***0.39*** | 0.006 | ***1.67*** | 0.06 | 4.0 | 2.21 |
| 120-A [d] | - | | - | | | 3.35 |
| 120-B [e] | 0.46 | 0.005 | 1.25 | 0.05 | | 0.27 |
| 121-A [d] | - | | - | | | 2.57 |
| 121-B [e] | 0.35 | 0.005 | 1.03 | 0.07 | | 0 |
| 127-A [d] | - | | - | | | 1.91 |
| 127-B [e] | 0.23 | 0.003 | 1.28 | 0.03 | | 0.59 |

[a] Exchange induced chemical shift differences in Hz between HSQC and HMQC spectra.
[b] Changes in cross peak intensity upon lowering the pH of F87E from 6.7 to 4.4 at 277K. Data taken from Figure 7 in ref.[4]
[c] Entries in bold italics indicate residues for which the sign of ΔωN could be determined from exchange induced chemical shift differences between HSQC and HMQC spectra.
[d] Changes in intensity for state A cross peaks upon lowering the pH of F87E from 6.7 to 4.4 at 277K. State A cross peaks do not exhibit relaxation dispersion.
[e] Fitted values of ΔωH and ΔωN for bound state B of F87E.
* Cross peaks that are overlapped or missing at 277K

Table S2. Values of the $^{13}$CO chemical shift difference ΔωC' between the ground and excited states determined by fitting the relaxation dispersion data shown in Figure S9. The data were fitted to the global $k_{ex}$ and $p_B$ derived from the $^{15}$N and $^{1}H_N$ dispersion profiles using the Carver-Richards equation.[3] Uncertainties were estimated from 200 Monte Carlo simulations. Residue numbers refer to the carbonyl that exhibits dispersion, detected via the amide cross peak of the i+1 residue that participates in the peptide bond.

| Residue | ΔωC' (ppm) | Error (ppm) |
|---|---|---|
| 11 | 0.87 | 0.10 |
| 12 | 0.73 | 0.11 |
| 18 | 1.05 | 0.06 |
| 19 | 0.91 | 0.10 |
| 20 | 0.83 | 0.07 |
| 22 | 0.83 | 0.09 |
| 53 | 0.74 | 0.11 |
| 54 | 0.83 | 0.11 |
| 60 | 0.71 | 0.16 |
| 61 | 0.70 | 0.15 |
| 62 | 1.58 | 0.17 |
| 65 | 1.45 | 0.12 |
| 66 | 0.93 | 0.08 |
| 67 | 0.72 | 0.14 |
| 73 | 0.86 | 0.10 |
| 74 | 0.90 | 0.08 |
| 75 | 0.75 | 0.08 |
| 77 | 0.81 | 0.08 |
| 80 | 0.83 | 0.10 |
| 92 | 0.81 | 0.10 |
| 93 | 2.40 | 0.12 |
| 105 | 1.16 | 0.09 |
| 119-B [a] | 2.25 | 0.10 |

[a] Fitted value of ΔωC' for T119 carbonyl (detected via the A120 amide) in state B of F87E.
